# Dual Inhibition of MYRF Cleavage by Its JM Region and PAN-1 CCT Gates Developmental Timing in *C. elegans*

**DOI:** 10.64898/2026.01.08.698494

**Authors:** Zhimin Xu, Xiaoting Feng, Haochen Lyu, Yong-Hong Yan, Yongqi Zhou, Zhizhi Wang, Fang Bai, Meng-Qiu Dong, Yingchuan B. Qi

## Abstract

Post-embryonic development proceeds through discrete stages, yet the mechanisms that coordinate transitions between stages remain incompletely defined. The transmembrane transcription factor MYRF undergoes intramolecular cleavage to release a nuclear-localized fragment essential for developmental progression in *C. elegans*. Previously, we showed that PAN-1 as a key partner required for MYRF activity during development. PAN-1 promotes MYRF trafficking to the cell membrane—an essential step for MYRF cleavage—through interactions between their extracellular domains (Xia et al., 2021). Here, we show that MYRF-1 cleavage and nuclear translocation oscillate with larval stage transitions in *C. elegans*, peaking mid-to-late stage and being suppressed during molts. Using endogenous gene editing and mutant reporters, we identify an uncharacterized juxtamembrane (JM) region in MYRF-1 that self-inhibits cleavage. JM deletion triggers premature MYRF-1 nuclear entry, early *lin-4* activation, growth defects, and adult lethality. We further demonstrate that the cytoplasmic tail (CCT) of the transmembrane protein PAN-1 acts as a predominant trans-inhibitor of MYRF-1 cleavage, coupling extracellular association with cytoplasmic inhibition. PAN-1 CCT deletion causes near-constitutive MYRF-1 nuclear accumulation, leading to premature *lin-4* activation, accelerated M-cell division, and larval lethality. Removing these inhibitory mechanisms on MYRF-1 cleavage overrides nutrient-responsive developmental checkpoints. These findings uncover dual inhibitory mechanisms governing MYRF-1 cleavage and establish MYRF-1 cleavage as a key gatekeeper of developmental timing.

## Introduction

MYRF (Myelin Regulatory Factor) is a unique transmembrane transcription factor that undergoes self-catalyzed cleavage at the membrane, releasing a transcriptionally active fragment that translocates to the nucleus^[1–4]^. Its founding member, mammalian MYRF, was first identified as a master regulator of oligodendrocyte terminal differentiation and postnatal myelination in the central nervous system^[5]^. Constitutive knockout of MYRF in mice results in early embryonic lethality^[5]^, underscoring its essential developmental functions beyond myelination. In humans, MYRF haploinsufficiency causes a spectrum of congenital disorders collectively termed MYRF-related Cardiac Urogenital Syndrome^[6]^, affecting the heart, lungs, and urogenital system, further highlighting its developmental importance^[1]^. Despite these insights, the roles of MYRF in embryonic development remain largely undefined. Moreover, whether MYRF cleavage is subject to regulation—and how such regulation contributes to its developmental functions—remains an open and intriguing question.

In *Caenorhabditis elegans*, MYRF is represented by two paralogous genes, *myrf-1* and *myrf-2*^[7]^. Functional studies have demonstrated that *myrf-1* is essential for postembryonic development^[7,8]^. It was first identified by Russell *et al.* as a partial loss-of-function allele that caused abnormal extra molting during the adult stage^[8]^. A stronger loss-of-function allele resulted in larval arrest with unshed cuticle during the L1–L2 molt, supporting a critical role for *myrf-1* in molting^[7,8]^. In an independent study of synaptic remodeling in DD neurons, Meng *et al.* identified *myrf-1(ju1121)* mutants that exhibited a block in synaptic remodeling and arrested in early larval stages^[7]^. Furthermore, *myrf-1* is required for activation of the heterochronic microRNA *lin-4*^[9]^, a key regulator of stage transitions^[10,11]^. In both DD synaptic remodeling and *lin-4* activation, *myrf-1* and *myrf-2* act cooperatively, with *myrf-1* playing a predominant role^[7,9]^. Together, these findings indicate that *myrf-1* coordinates multiple developmental programs—including molting, synaptic remodeling, and stage-timing regulation—during early larval development.

As outlined above, a distinctive feature of MYRF is its ability to undergo auto-cleavage, allowing a fragment to break away from the membrane-anchored precursor and function as a nuclear transcription factor^[3,4,12,13]^. In mammalian cells, full-length MYRF localizes to the endoplasmic reticulum (ER), where it self-cleaves through a conserved intramolecular chaperone auto-processing (ICA) domain. This domain, structurally related to the intramolecular chaperone domains of bacteriophage tailspike proteins, forms a trimeric complex that catalyzes an internal peptide bond cleavage via a serine–lysine catalytic dyad^[14]^. Cleavage of MYRF releases the N-terminal fragment containing the DNA-binding and transcriptional activation domains, which then translocates to the nucleus to regulate gene expression.

This self-cleavage mechanism of MYRF is conserved in *C. elegans*^[7]^. Both MYRF paralogs, *myrf-1* and *myrf-2*, undergo cleavage to generate a nuclear-localized N-terminal fragment^[7]^. Functional rescue experiments demonstrate that both cleavage and nuclear translocation of N-MYRF are essential for MYRF-1 activity. Furthermore, endogenous gene editing that mutates key residues within the ICA domain cleavage site—*myrf-1(syb1487[S483A, K488A])*—blocks cleavage and causes MYRF-1 to remain at the cell membrane^[15]^. The *myrf-1(syb1487)* mutants exhibit developmental arrest at the end of the L1 stage, phenocopying *myrf-1* null mutants. Conversely, overexpression of N-MYRF-1 is sufficient to drive premature synaptic rewiring in DD neurons^[7]^. However, for endogenous function, N-MYRF must assemble into a trimeric form. Gene-edited mutants expressing only the N-terminal fragment—*myrf-1(syb1491[1–482])*—presumably as a monomer, display null-like defects despite robust nuclear localization of MYRF-1[1–482]^[15]^.

While the cleavage process of MYRF is conserved, two key discrepancies exist between mammalian and *C. elegans* MYRF proteins: their subcellular localization prior to cleavage and whether cleavage occurs constitutively. When expressed in cultured cells, full-length mammalian MYRF localizes to the endoplasmic reticulum (ER) and undergoes constitutive self-cleavage^[2]^. In contrast, recent studies in *C. elegans* indicate that full-length MYRF traffics to the cell membrane, where its cleavage is temporally controlled—active in late L1 but minimal in early and mid-L1 stages^[15]^. The trafficking of *C. elegans* MYRF from the ER to the cell membrane depends on its binding partner PAN-1, a transmembrane protein with a leucine-rich repeat (LRR) domain^[15]^. The interaction between MYRF-1 and PAN-1 occurs via their exoplasmic regions^[15]^. In *pan-1* null mutants (*gk142*), MYRF-1 fails to reach the cell membrane and is degraded in the cytoplasm, likely due to instability of its exoplasmic portion^[15]^. Deletion of the exoplasmic region in *myrf-1(syb1313)* mutants causes the protein to be stably retained in the cytoplasm but poorly cleaved^[15]^. Furthermore, even modest overexpression of *C. elegans* MYRF-1 leads to its accumulation in ER-like structures without processing^[8]^. Together, these observations suggest that MYRF cleavage in *C. elegans* is not constitutive but rather regulated by additional factors that remain to be identified.

Despite these insights, the mechanisms that regulate the timing of MYRF cleavage remain unresolved, and the physiological significance of its temporal control is unclear. In this study, we set out to dissect the regulatory control of MYRF-1 cleavage in *C. elegans* and identified two inhibitory modules—one intrinsic to MYRF-1 and another mediated by the transmembrane protein PAN-1, with the latter exerting a predominant effect. This finding uncovers a second functional role of PAN-1 beyond facilitating MYRF-1 trafficking. Disruption of these inhibitory mechanisms leads to premature nuclear accumulation of N-MYRF-1, resulting in early activation of next-stage developmental programs and bypass of starvation-induced developmental checkpoints. Together, these results establish MYRF-1 cleavage as a key regulatory node that coordinates stage progression.

## Results

### MYRF-1 Cleavage-Nuclear-translocation Oscillates During Each Larval Stage

We previously inserted a GFP tag into the N-terminal region of the *myrf-1* gene, generating the allele *ybq14[gfp::A172]*, in which GFP marks both full-length MYRF-1 and the cleaved N-terminal fragment (N-MYRF-1)^[7]^. In early to mid-L1 larvae, GFP was localized to the cell membrane and absent from the nucleus^[15]^ (Figure 1A-C). By late L1, however, nuclear GFP became enriched while membrane signals diminished, indicating increased MYRF-1 cleavage and nuclear translocation (Figure 1A-C). Although *myrf-1* transcript levels peak at late L1^[16]^, the increase in N-MYRF-1 is unlikely due to transcript abundance alone, as overexpression of *myrf-1* does not effectively increase N-MYRF-1 levels^[7,8]^ (Figure 1-figure supplement 1A). Western blot analysis further supports temporally regulated cleavage: the cleaved N-MYRF-1 fragment is detectable in synchronized late L1 (14 h) but not in early L1 (6 h) animals (Figure 1D).

**Figure 1.**
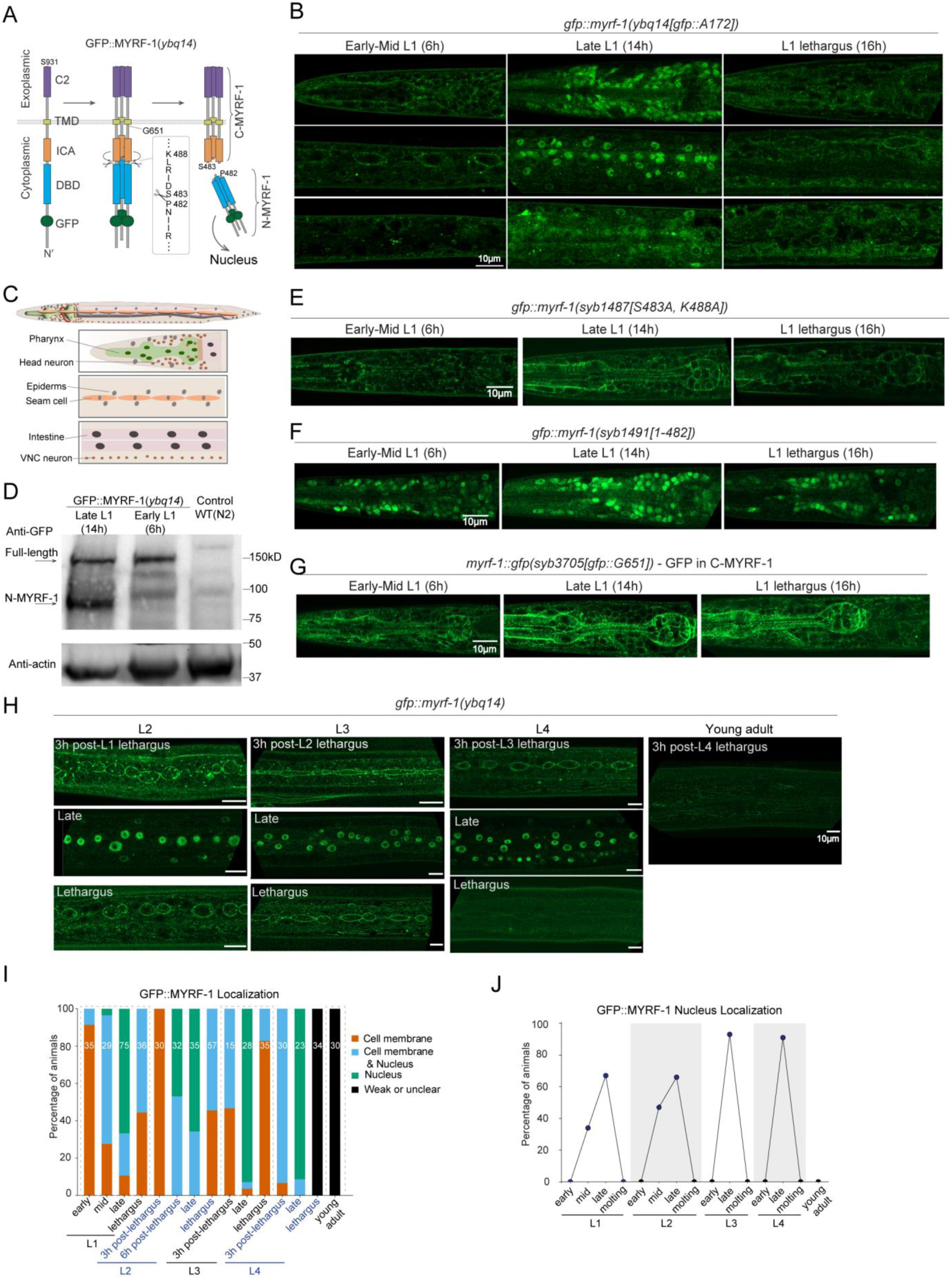
Developmentally Regulated Cleavage and Localization of MYRF-1. (A) Domain structure and cleavage of endogenously tagged MYRF-1. GFP was inserted at Ala172 (*ybq14* allele) or mEGFP at Gly651 (*syb3705* allele). Self-cleavage is mediated by the trimerized ICA domain, with key catalytic residues S483 and K488, producing an N-terminal fragment (1–P482). (B) GFP::MYRF-1(*ybq14*) localization during L1–L2 transition correlated with seam cell divisions, visualized with Pscm::RFP::PH(*ybqSi296*). Seam cells categorized by division/fusion state: Pre-1st division; 1st division only; 1st and 2nd divisions; Post-2nd division (prior to fusion); and Fusion/Elongation (elongated daughter cells). Right: quantification of MYRF-1 localization in each category. (C) Stage-specific GFP::MYRF-1(*ybq14*) localization in L1 larvae. Localization shifts from membrane-only (early L1) to nuclear (mid/late L1), and back to membrane during lethargus. Lethargus was defined by cessation of pharyngeal pumping. Three sagittal sections shown: mid-head (pharynx, head neurons), lateral trunk (epidermis, seam), and mid-trunk (intestine, ventral nerve cord). See (C) for schematic. (D) Schematic of major cell types visible in (B) images. Top: head region (pharynx, neurons); middle: lateral trunk (epidermis, seam); bottom: midline trunk (intestine, VNC). Muscle and gonad not shown. (E) Western blot detection of MYRF-1 cleavage. Protein extracts from early (6 h) and late (14 h) L1 larvae of GFP::MYRF-1(*ybq14*) and N2 controls. Cleaved N-terminal (81 kDa) band appears only in late L1; full-length (131 kDa) detected in both. (F) Cleavage-defective mutant *gfp::myrf-1(syb1487[S483A, K488A])*. GFP signal remains membrane-localized throughout L1, with no nuclear enrichment. (G) N-terminal only mutant *gfp::myrf-1(syb1491[1–482])*. GFP is consistently nuclear, with no membrane signal. (H) C-terminal tagged GFP in *gfp::myrf-1(syb3705[GFP::G651])*. GFP signal is detected on the membrane throughout L1. (I) Oscillatory MYRF-1 localization from L2 to adulthood. GFP::MYRF-1 shows membrane-localized signal in early stages, nuclear signal in mid/late stages, and membrane-signal again during lethargus. Signal disappears in young adults. Early larval stages were defined as 3 h post-lethargus; late L2, L3, and L4 as 25 h, 35 h, and 44 h, respectively. (J) Quantification of GFP::MYRF-1 subcellular localization. (K) Percentage of nuclear-enriched GFP::MYRF-1 at each stage.

To confirm that cleavage produces the nuclear MYRF-1 fragment, we examined two previously characterized *myrf-1* variants^[15]^. In *myrf-1(syb1487[S483A, K488A])*, the cleavage sites are mutated, preventing processing, and GFP remains at the membrane throughout L1 (Figure 1E; Figure 1-figure supplement 1B). In contrast, *myrf-1(syb1491[1–482])* expresses only the N-terminal portion due to an in-frame deletion of the C-terminal region; this results in constitutive nuclear GFP localization at all stages (Figure 1F; Figure 1-figure supplement 1C). We also generated a new allele, *myrf-1*(*syb3705[GFP::G651])*, in which GFP is inserted at G651 to tag both the full-length and C-terminal MYRF-1 fragments. In this allele, GFP remains localized to the membrane at all stages (Figure 1G; Figure 1-figure supplement 1D). Together, these results demonstrate that MYRF-1 undergoes regulated intramolecular cleavage, with cleavage activity increasing during late L1 to generate the nuclear-localized active form.

In larvae beyond the L1 stage, we previously observed a heterogeneous pattern of MYRF-1 localization: some individuals exhibited predominantly nuclear signals, others showed both membrane and nuclear localization, and some displayed only membrane-associated signals^[15]^. This variability suggests that MYRF-1 cleavage does not switch into a constitutive “on” state after late L1. Through a more detailed analysis of synchronously staged animals, we found that nuclear N-MYRF-1 signals declined specifically during the L1–L2 molting period (Figure 1B), defined here as the developmental transition marked by behavioral quiescence (lethargus) and ecdysis. As larvae entered lethargus, nuclear GFP signals gradually diminished and became undetectable, while faint signals reappeared at the membrane (Figure 1B). Over the next three hours, as animals completed molting and transitioned into L2, GFP::MYRF-1 was exclusively detected at the membrane (Figure 1H). Cleavage activity resumed in mid-to-late L2, indicated by renewed nuclear GFP::MYRF-1 and the corresponding loss of membrane signal at peak cleavage (Figure 1H). A similar pattern of cleavage suppression was observed during L2 molting (Figure 1H-J). This membrane-to-nucleus cycle repeated throughout the L3 and L4 stages (Figure 1H-J). By the L4–adult molt and in adults, GFP::MYRF-1 signals became barely detectable (Figure 1H-J). These results indicate that MYRF cleavage activity follows an oscillatory pattern—suppressed during the early phase of each larval stage, activated during the mid-to-late phase, and suppressed again during molting—until MYRF expression is inactivated at the L4–adult transition. MYRF is broadly expressed throughout the body (excluding germ cells), and the membrane– nuclear localization pattern is generally consistent across tissues, suggesting that MYRF cleavage is regulated by a systemically controlled temporal mechanism.

### MYRF-1 Cleavage Is Inhibited During Starvation-Induced Developmental Arrest

Nutrient-associated developmental checkpoints at early L3 and early L4 have been well characterized and are induced by starvation during the late phase of the preceding stage (late L2 and late L3, respectively)^[17]^. Under these conditions, animals enter a systemic developmental arrest mediated by insulin and steroid hormone signaling pathways. To test whether such nutritional fluctuations influence MYRF-1 cleavage, we transferred late L1 animals to fresh plates devoid of food. After 8 hours of starvation, GFP::MYRF-1 was exclusively localized to the cell membrane, with no detectable nuclear signal (Figure 2A). To assess developmental progression, we used seam cell division patterns as morphological landmarks, visualized by an RFP::PH membrane marker in a GFP::MYRF-1 background (Figure 2B). In well-fed animals, MYRF-1 remained membrane-localized during the first and second seam cell divisions, whereas nuclear MYRF-1 signals emerged only after completion of the second division and peaked during epidermal fusion and elongation of seam daughter cells (Figure 2B). In contrast, most late-L1-starved animals arrested between the first and second seam cell divisions, exhibited rounded, non-elongated seam cells, and showed exclusively membrane-localized MYRF-1 (Figure 2C). These observations indicate arrest at early L2 and support the existence of a developmental checkpoint at this stage.

**Figure 2.**
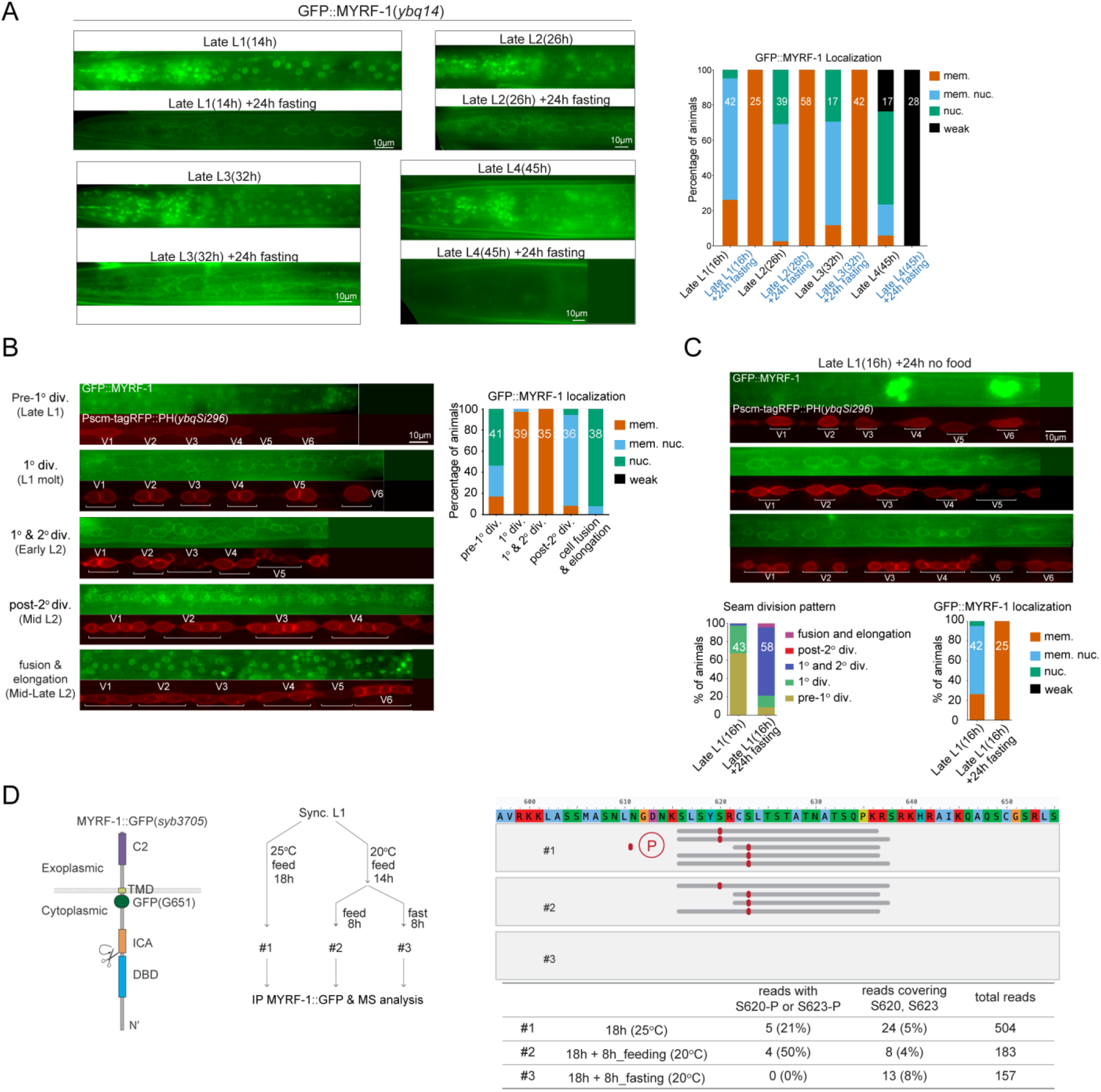
MYRF-1 Cleavage Is Suppressed at Nutrient-Dependent Developmental Checkpoints. (A) GFP::MYRF-1 localization under starvation initiated at late L1, L2, L3, or L4. In all groups, GFP remains membrane-associated, except in late L4-starved animals, where signal is lost. Right: Quantification of GFP localization. (B) GFP::MYRF-1 localization is altered by nutritional status. In late L1, it is enriched in the nucleus under feeding conditions. After 8 hours of food deprivation starting in late L1, GFP::MYRF-1 becomes localized to the cell membrane, while control animals that continue feeding show high nuclear localization. (C) Representative GFP::MYRF-1 localization and seam cell pattern 24 hours after late L1 food removal. Below: Quantification of seam cell categories from (B); Quantification of GFP::MYRF-1 localization patterns in late L1 starvation assay. (D) IP–MS analysis of MYRF-1 under nutrient-rich and food-deprived conditions using *myrf-1::gfp (syb3705)*. Left: Schematic of the C-terminal GFP fusion at Gly651. Middle: Experimental design for culture and starvation treatment. Right: Mass-spectrometry reads covering residues S620 or S623 that contain identified phosphorylation in well-fed versus food-deprived samples. Only phosphopeptide-containing reads are shown. Bottom table: Percentage of phosphopeptide reads among those covering S620/S623, and the proportion of reads covering these residues relative to total MYRF-1 reads.

We next examined MYRF-1 localization following food removal at later stages. When food was removed during late L2 or late L3, GFP::MYRF-1 remained membrane-associated, and seam cells displayed rounded morphologies consistent with early-stage arrest (Figure 2A), as previously reported. When food was removed during late L4, GFP::MYRF-1 signals became undetectable (Figure 2A), consistent with developmental downregulation of MYRF at this stage. Together, these results demonstrate that nutrient deprivation during the late phase of a larval stage induces arrest at the early phase of the subsequent stage, during which MYRF-1 cleavage is consistently suppressed.

We hypothesized that contrasting nutrient-rich and nutrient-deprived conditions might be exploited to identify factors associated with MYRF-1 cleavage activation or inhibition. To this end, we performed immunoprecipitation using a C-terminally tagged allele, *myrf-1(syb3705[GFP::G651])*, from whole-animal extracts prepared from well-fed animals or animals subjected to 8 hours of starvation beginning at late L1 (Figure 2D). Co-immunoprecipitated proteins were analyzed by mass spectrometry. This IP–MS analysis identified factors differentially associated with MYRF-1 under nutrient-rich versus nutrient-deprived conditions and revealed phosphorylation at residues S620 and S623 of MYRF-1 specifically in nutrient-rich samples (Figure 2D). These phosphosites were not detected in nutrient-deprived samples, despite comparable overall MYRF-1 coverage (Figure 2D; Figure 2—figure supplement 1A). To explore the functional relevance of this modification, we generated a gene-edited GFP::MYRF-1[S623A] allele to mimic a non-phosphorylatable state. This mutant showed no overt developmental defects, although a modest trend toward reduced nuclear MYRF-1 enrichment was observed (Figure 2—figure supplement 1B).

Although these data did not yield a definitive mechanistic interpretation for MYRF-1 phosphorylation, they directed our attention to an uncharacterized juxtamembrane (JM) region of MYRF-1, spanning approximately 60 amino acids (A598–S655) between the transmembrane domain and the ICA domain (Figure 3A). Structural predictions using AlphaFold indicate that the JM region lacks a defined fold^[18]^, consistent with an intrinsically flexible segment. Such regions often function as regulatory interfaces, and the JM region lies immediately adjacent to the α-helical segment of the ICA domain that is critical for trimerization. We therefore hypothesize that the JM region may influence accessibility of this α-helix and thereby modulate MYRF-1 trimerization and cleavage.

**Figure 3.**
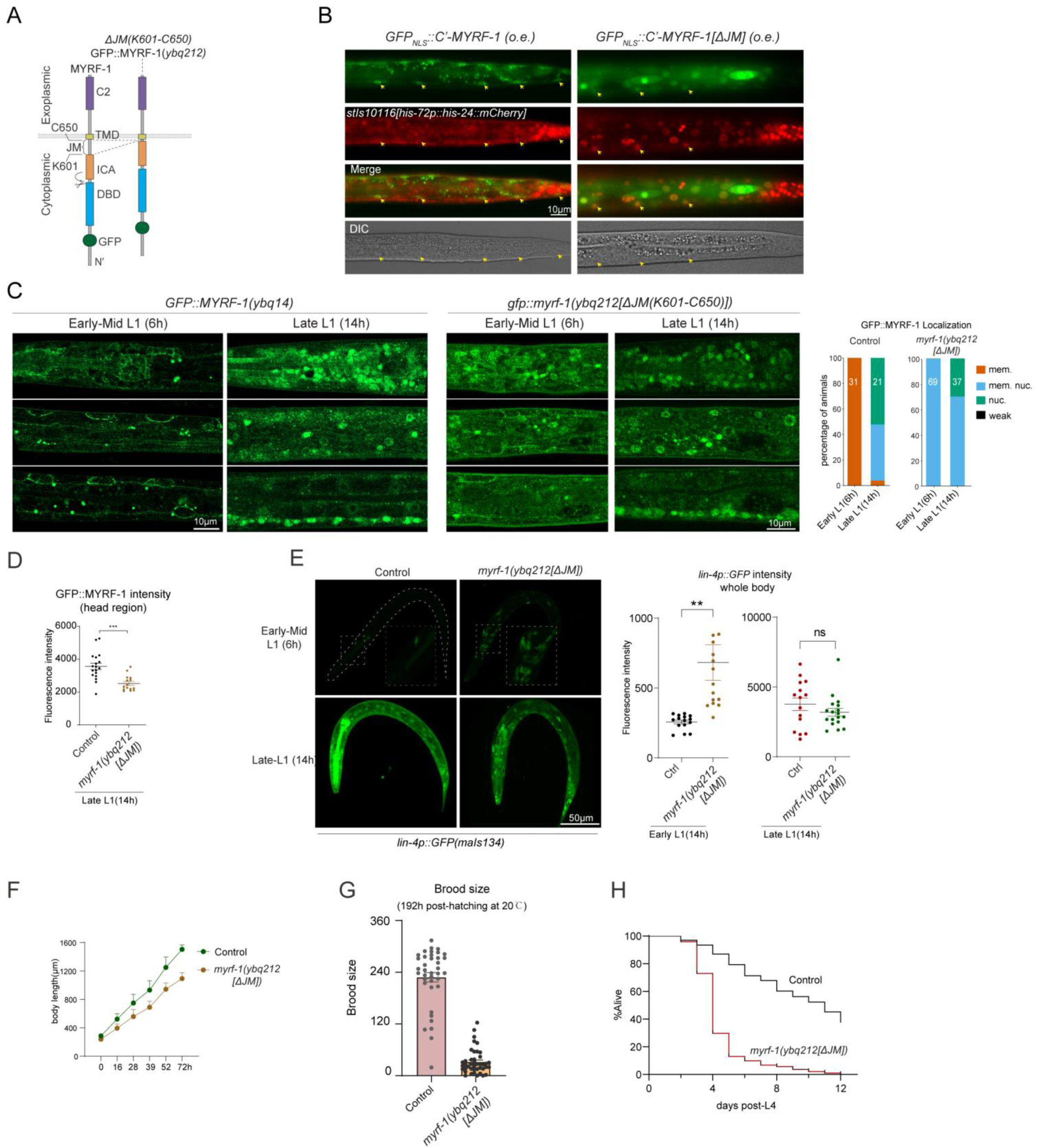
The Juxtamembrane Region of MYRF-1 Inhibits Cleavage and Regulates Development. (A) Schematic of *gfp::myrf-1(ybq212[ΔJM])* allele, with deletion of the juxtamembrane (JM) region (K601–C650). (B) GFP localization upon overexpression of NLS::GFP::MYRF-1[G477-C’]. GFP remains cytoplasmic with intact JM; nuclear enrichment is observed when the JM region (K601–C650) is deleted. Nuclei marked by his-24::mCherry. (C) Subcellular localization of GFP::MYRF-1[ΔJM]. GFP is detected at both the membrane and nucleus in early-mid L1 (6 h), with increased nuclear accumulation by late L1 (14 h). Right: Quantification of GFP localization in wild-type and *myrf-1[ΔJM]* animals at early-mid and late L1 stages. (D) Reduced GFP intensity in *myrf-1[ΔJM]* mutants. Mean signal intensity in the head region ± SEM; ***p < 0.001, t-test. (E) *lin-4p::gfp* reporter expression in *myrf-1[ΔJM]* mutants vs. controls. Reporter activated precociously in mutants at early late L1. Right: quantification of GFP intensity (mean ± SEM; ***p < 0.001, t-test). (F) Body length over time in *myrf-1[ΔJM]* vs. wild-type. Mean ± SD from L1 to young adult (n = 25 per genotype per stage. (G) Total progeny counts (± SEM) of wild-type and *myrf-1[ΔJM]* at 192 h post-hatching; each data point represents progeny count from an individual animal. (H) Lifespan analysis of *myrf-1[ΔJM]* vs. wild-type. Kaplan-Meier survival curve monitored from day 1 post-L4 (n ≥ 200 per genotype).

### MYRF Cytoplasmic Juxtamembrane (JM) Region Self-Inhibits Cleavage

To test MYRF-1 JM’s function, we deleted the JM region from a transgenic construct in which the N-terminal MYRF-1 was replaced with NLS::GFP to minimize the toxicity associated with MYRF-1 overexpression (Figure 3B). Despite this precaution, overexpression of the C-terminal variant still caused lethality, and we obtained only a few live transgenic progenies showing sparse GFP expression. In these animals, the control construct with an intact JM region localized predominantly to the cytoplasm (Figure 3B), consistent with our recurrent observation that elevated MYRF-1 expression does not enhance cleavage efficiency. In contrast, deletion of the JM region resulted in strong nuclear GFP localization (Figure 3B), revealing a cleavage-inhibitory role for the JM region.

To examine the JM region under endogenous conditions, we deleted it (K601–C650) by genome editing, generating the *gfp::myrf-1(ybq212[ΔJM(K601–C650)])* mutant (Figure 3A, C). At early L1, GFP signals were detected in the nuclei of a subset of cells in ΔJM mutants, in contrast to the absence of nuclear signals in wild-type animals (Figure 3C). Notably, GFP::MYRF-1[ΔJM] remained strongly membrane-localized at this stage, indicating that cleavage activation was only partial—less efficient than the robust cleavage observed in overexpression experiments. Additionally, the overall GFP fluorescence intensity was reduced in *myrf-1[ΔJM]* mutants (Figure 3D), suggesting that the JM region also contributes to MYRF-1 protein stability. Together, these findings demonstrate that the JM region functions as an intrinsic inhibitory element that restricts MYRF-1 cleavage, and its removal partially releases MYRF from this control.

### MYRF-1[ΔJM] Mutants Display Mild Precocity and Reduced Viability

We previously reported that transcriptional activation of the *lin-4* microRNA critically depends on MYRF cleavage^[9]^. To assess whether deletion of the JM region affects this process, we examined expression of the *lin-4* transcriptional reporter *lin-4p::gfp(maIs134)* in *myrf-1[ΔJM]* mutants (Figure 3E). At 6 hours post-hatching—a stage when *lin-4* is not yet activated in control animals—we observed premature reporter expression in the mutants. This precocious activation was confirmed using the endogenous *lin-4* transcriptional reporter *umn84* (Figure 5—figure supplement 1C). Notably, reporter expression was restricted to a subset of cells, rather than being widespread throughout the body, and reporter brightness closely correlated with nuclear accumulation of GFP::MYRF-1. These findings support that deletion of the JM region elevates MYRF cleavage activity, leading to early *lin-4* activation in a spatially restricted manner.

We next assessed the developmental and physiological consequences of JM deletion in *myrf-1(ybq212[ΔJM])* mutants. Although these mutants developed slightly slower than wild-type animals, nearly all progressed to the gravid adult stage (Figure 3—figure supplement 1A). However, they remained visibly smaller than controls throughout both larval and adult stages (Figure 3F), and were inactive or slow-moving, suggesting potential neuromuscular or neural defects. While fertile, *myrf-1[ΔJM]* mutants produced significantly fewer progeny (10–50 per animal) compared to wild type (200–300) (Figure 3G). Most strikingly, they exhibited highly penetrant lethality during early adulthood, typically dying between 2 and 4 days of adult life (Figure 3H). The lethality may be attributed to bagging, a common phenotype in mutants (Figure 3—figure supplement 1A); however, the exact defects require further investigation. Together, these findings demonstrate that the JM region of MYRF-1 plays a critical role in regulating cleavage timing and ensuring proper growth, fertility, and adult survival.

### MYRF-1[ΔJM] Partially Bypasses Late-Stage Nutrient Checkpoints

We examined whether MYRF-1 cleavage inhibition contributes to nutrient checkpoints. Upon starvation at late L1, both *myrf-1(ΔJM)* mutant and control animals showed rounded, non-elongated seam cells^[19]^, indicating no bypass of early L2 checkpoint (Figure 4A). However, in late L2 starvation assays, *myrf-1*[ΔJM] mutants exhibited one or two rounds of SM divisions, whereas control animals showed none (Figure 4B), suggesting a partial bypass of the early L3 checkpoint, despite the absence of (L3) molting behavior. In late L3 starvation assays, many *myrf-1*[ΔJM] mutants, although arrested with early L4–like body size, developed visible vulvae and underwent ecdysis—features not observed in controls (Figure 4C). These results indicate that while *myrf-1*[ΔJM] mutants fail to overcome the early L2 nutrient checkpoint, they can partially bypass later-stage checkpoints under nutrient-restricted conditions.

**Figure 4.**
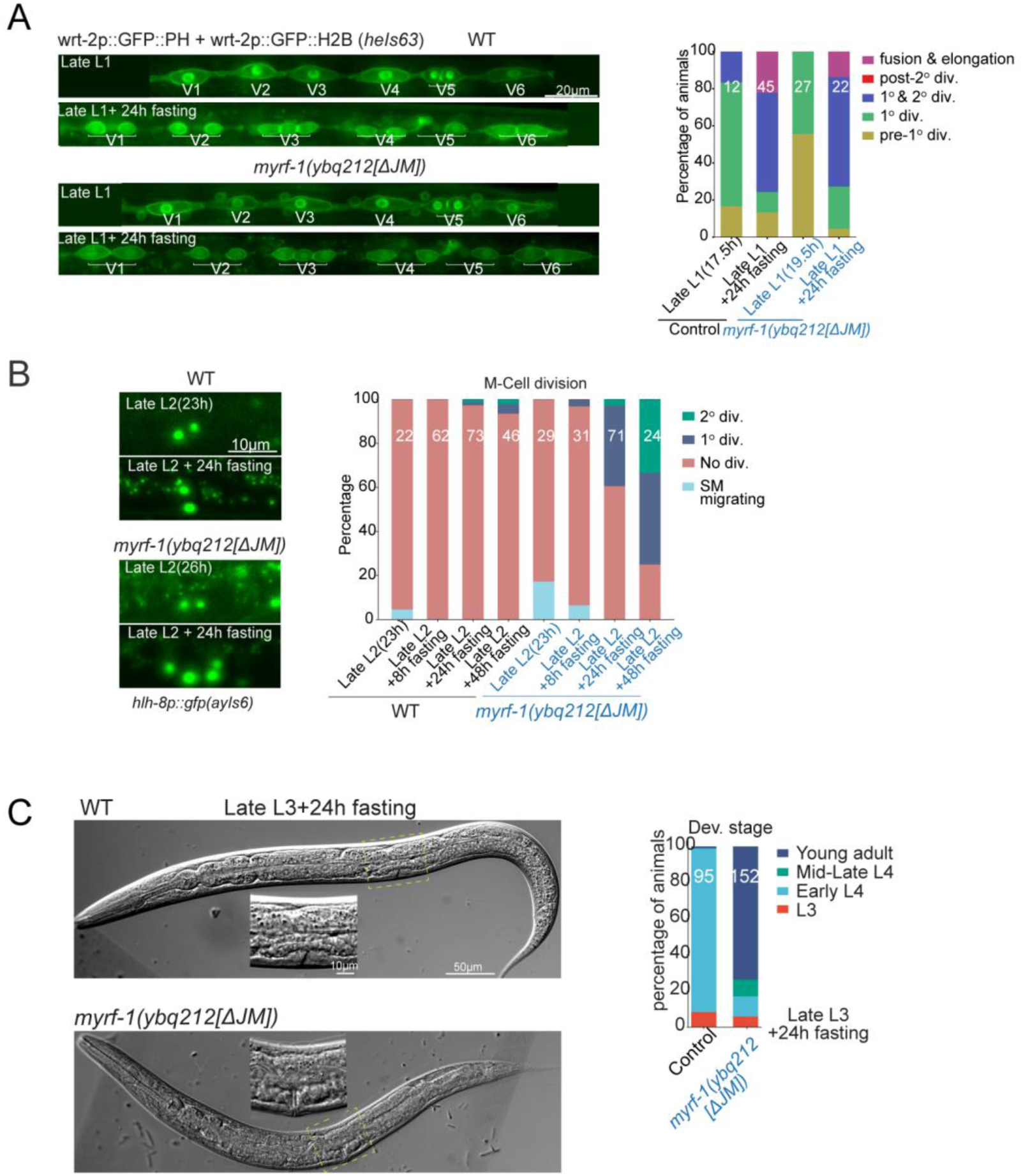
MYRF-1[ΔJM] Partially Bypasses Late-Stage Nutrient Checkpoints. (A) Seam cell divisions (marked by *heIs63*) 24 h after late L1 starvation. *myrf-1[ΔJM]* mutants resemble controls, with most seam cells rounded and between the first and second divisions. Right: Quantification of seam cell categories. (B) Sex myoblast division assessed using *hlh-8p::gfp* at late L2 and 24 h post-food removal. *myrf-1[ΔJM]* mutants show advanced divisions. Right: quantification at late L2; 8h, 24h and 48h starvation. (C) Vulval development 24 h after late L3 starvation. *myrf-1[ΔJM]* mutants show adult vulvae and cuticle shedding, while controls remain in early L4. Right: stage quantification based on vulva morphology.

### PAN-1 Cytoplasmic Tail Predominantly Inhibits MYRF-1 Cleavage

Given the more efficient cleavage observed in the overexpressed MYRF-1[ΔJM] transgene compared to the only mildly premature cleavage seen in the *myrf-1[ΔJM]* mutant, we hypothesized that an additional factor contributes to MYRF cleavage regulation. This putative factor is likely limiting in quantity and, under physiological conditions, interacts with MYRF to inhibit its cleavage. While exploring potential trans-acting inhibitors, we considered PAN-1, a consistent binding partner of MYRF-1. Our previous work showed that PAN-1 and MYRF-1 interact via their exoplasmic domains^[15]^. However, the cytoplasmic tail (CCT) of PAN-1 is well-positioned to access the cytoplasmic ICA domain of MYRF-1, raising the possibility that it could directly influence cleavage activity (Figure 5A). The PAN-1 CCT comprises 55 amino acids (K539–F594) and lacks predicted secondary structure, suggesting a potentially flexible regulatory role.

**Figure 5.**
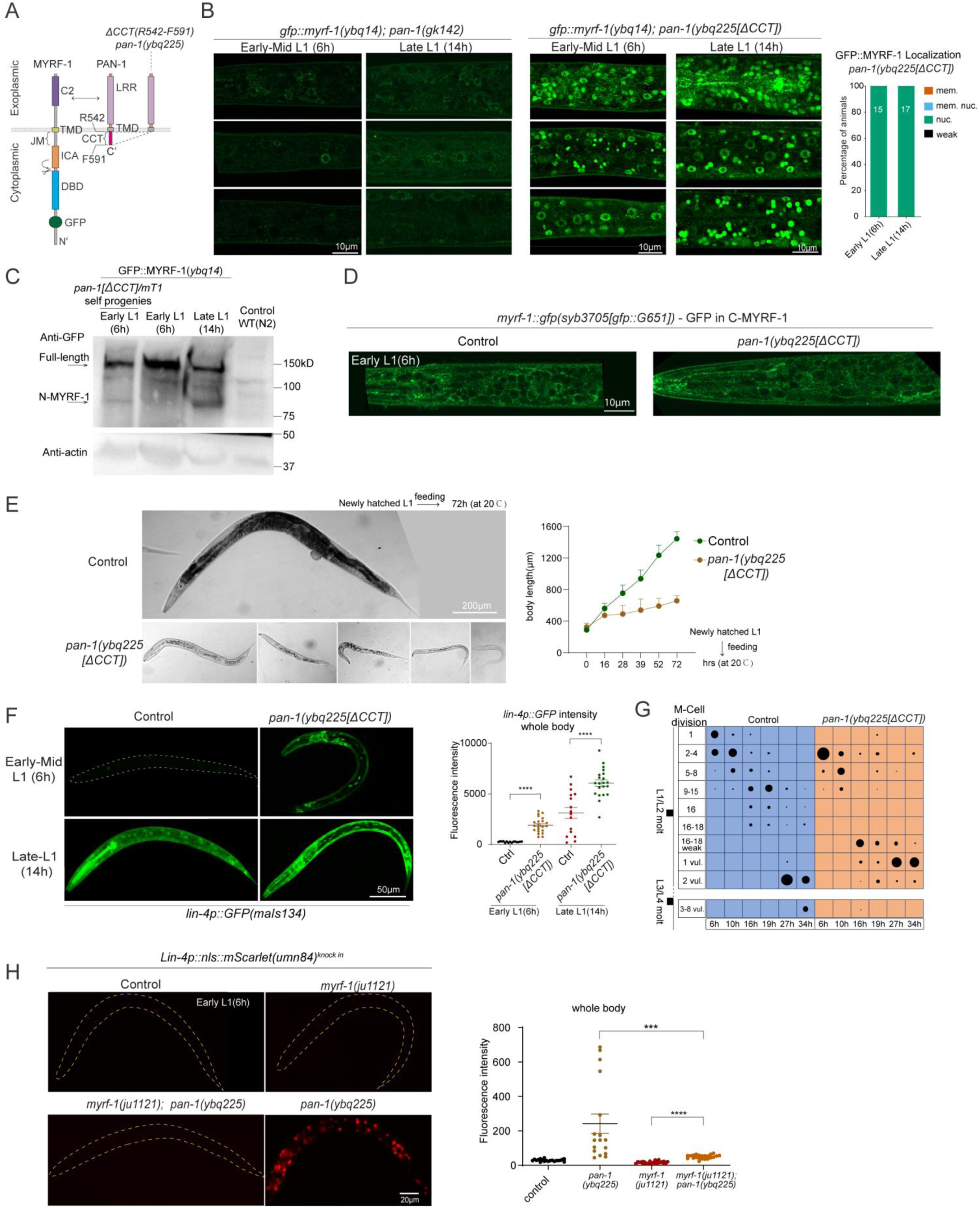
PAN-1 CCT Inhibits MYRF-1 Cleavage and Controls Developmental Progression. (A) Schematic of the *pan-1(ybq225[ΔCCT])* allele with deletion of the cytoplasmic tail (CCT); also shows extracellular interaction between PAN-1 and MYRF-1. (B) GFP::MYRF-1(*ybq14*) localization in *pan-1* mutants: in *pan-1(gk142)* null mutants MYRF-1 is degraded in the cytoplasm; in *pan-1[ΔCCT]* mutants GFP::MYRF-1 is strongly nuclear at 6 h and 14 h. Right: quantification of GFP localization. (C) Western blot analysis of MYRF-1 cleavage in *pan-1[ΔCCT]* mutants; extracts from *gfp::myrf-1*(*ybq14*); *pan-1[ΔCCT]/mT1* progeny show an 81 kDa cleaved N-terminal MYRF-1 band (∼10% homozygous mutants). (D) C-terminal GFP fusion *myrf-1::gfp(syb3705)* in *pan-1[ΔCCT]* shows membrane localization at early L1, indicating intact trafficking. (E) Developmental arrest in *pan-1[ΔCCT]* mutants: Left: representative animals at 3 days post-hatching showing arrest between L1–L3; Right: body length over time (mean ± SD). (F) *lin-4p::gfp* expression is prematurely activated in *pan-1[ΔCCT]* at 6 h and 14 h. Right: quantification (mean ± SEM; ****p < 0.0001, t-test). (G) M-cell lineage progression assessed with *hlh-8p::gfp(ayIs6)*; classification based on division and migration status; dot size reflects population proportion. (H) Endogenous *lin-4* reporter (*umn84*) expression in control, *myrf-1(ju1121)*, *pan-1[ΔCCT]*, and *myrf-1; pan-1[ΔCCT]* double mutants at 6 h; premature *lin-4* activation in *pan-1[ΔCCT]* is suppressed in the double mutant. Right: Quantification of reporter intensity (mean ± SEM; ***p < 0.001, ****p < 0.0001; t-test).

To test this, we generated a *pan-1* CCT deletion mutant (*pan-1(ybq225[ΔCCT])*), removing residues R542–F591 (Figure 5A, B). In this background, GFP::MYRF-1 displayed strong nuclear localization in most cells at both early and late L1 stages (Figure 5B), in sharp contrast to *pan-1* null mutants, in which MYRF-1 fails to traffic to the membrane and is largely degraded in the cytoplasm^[15]^ (Figure 5B). Western blot analysis further confirmed premature cleavage in *pan-1[ΔCCT]* mutants: a band corresponding to the N-terminal fragment of MYRF-1 was detected, whereas this band was minimal in wild-type samples (Figure 5C) (see Methods for genotype details).

To further examine cleavage processing, we analyzed the MYRF-1(*syb3705*[GFP::G651]) allele, which tags both full-length MYRF-1 and the post-cleavage C-MYRF-1. In *pan-1[ΔCCT]* mutants, GFP signals remained consistently at the cell membrane across developmental stages (Figure 5D; Figure 5-figure supplement 1A), indicating that full-length MYRF-1 is properly trafficked and that cleavage occurs at the membrane. These results suggest that the exoplasmic PAN-1–MYRF-1 interaction remains intact in the ΔCCT mutant, and that the PAN-1 with the CCT deleted still supports MYRF-1 membrane localization. Taken together, our findings reveal a dual role for PAN-1: its exoplasmic LRR domain facilitates MYRF-1 trafficking, while its cytoplasmic CCT acts as a critical inhibitory module that suppresses MYRF cleavage.

### PAN-1[ΔCCT] Causes Pronounced Developmental Precocity and Larval Lethality

Deletion of the PAN-1 CCT domain resulted in larval lethality (Figure 5E; Figure 5—figure supplement 1B). The *pan-1[ΔCCT]* mutants exhibited variable developmental arrest ranging from the L1 to L3 stages based on body size, but rarely progressed to L4. For comparison, *pan-1* null mutants (*gk142*) also showed early larval arrest; however, their arrest was more uniform, and the animals survived for at least over one week without immediate lethality^[20]^. In contrast, *pan-1[ΔCCT]* mutants died shortly after arrest, indicating distinct phenotypic consequences despite the shared early arrest phenotype.

We previously reported that loss of *pan-1* leads to MYRF-1 trafficking failure and degradation^[15]^ (Figure 5B), with a complete loss of *lin-4* microRNA transcription^[9]^. In contrast, the *pan-1[ΔCCT]* mutant exhibited strong precocious *lin-4* activation, as revealed by the *lin-4p::gfp(maIs134)* reporter (Figure 5F) and the endogenous transcriptional reporter *umn84* (Figure 5—figure supplement 1C). By 6 hours post-hatching, many cells in *pan-1[ΔCCT]* animals expressed *lin-4*, whereas wild-type animals did not activate expression until at least 10 hours. By 14 hours, *lin-4* expression was significantly elevated in *pan-1[ΔCCT]* compared to controls (Figure 5F). This premature *lin-4* activation is consistent with the uncontrolled MYRF-1 cleavage observed in *pan-1[ΔCCT]* mutants. Importantly, this effect was dependent on MYRF-1 activity: in the *myrf-1(ju1121); pan-1[ΔCCT]* double mutant, *lin-4* was no longer prematurely activated, or not activated at all (Figure 5H), confirming that MYRF-1 functions downstream of PAN-1 CCT to mediate this precocious activation.

To evaluate developmental progression, we examined M-cell division using the *hlh-8p::GFP* reporter, which marks the M cell and its stem-like descendants. In *pan-1[ΔCCT]* mutants, M-cell division occurred earlier than in controls, with a rapid succession of divisions between 6 and 16 hours (Figure 5G; Figure 5—figure supplement 1D), indicating accelerated L1 development. However, a prolonged period of weak *hlh-8p::GFP* expression followed before signal reappeared in the remaining stem cells—the sex myoblasts—suggesting delayed specification (Figure 5G). Migration of these cells to the presumptive vulval region was also delayed compared to controls (Figure 5G). These results indicate that *pan-1[ΔCCT]* mutants undergo premature L1 development, but the progression does not extend robustly beyond late L1, likely leading to arrested and lethal outcomes due to misaligned developmental timing.

We next assessed the genetic interaction between the two inhibitory modules: PAN-1 CCT and MYRF-1 JM. In the *gfp::myrf-1[ΔJM]; pan-1[ΔCCT]* double mutant, GFP localization was predominantly nuclear, with only faint membrane signals (Figure 5—figure supplement 1E), similar to the *pan-1[ΔCCT]* single mutant. However, the double mutants exhibited more severe arrest than either single mutant (Figure 5—figure supplement 1B), with most animals arrested at the L1–L2 transition. These findings suggest that the PAN-1 CCT and MYRF-1 JM regions contribute overlapping as well as partially independent regulatory influences on MYRF cleavage and developmental timing.

### Loss of MYRF-1 Cleavage Inhibition Disrupts Starvation-Induced L1 Arrest

When newly hatched L1 larvae encounter the absence of food (Figure 6A), they enter a facultative diapause, halting all post-embryonic development, including growth and cell division^[21]^. We investigated whether MYRF-1 cleavage regulation plays a role in L1 diapause. Under food-deprived conditions, GFP::MYRF-1 signals remained exclusively membrane-localized throughout the extended L1 diapause (Figure 6B), indicating suppressed MYRF-1 cleavage in alignment with arrested development. In contrast, *pan-1[ΔCCT]* mutants showed predominant nuclear localization of GFP::MYRF-1 (Figure 6B), suggesting that PAN-1’s CCT domain is essential for repressing MYRF-1 cleavage during diapause.

**Figure 6.**
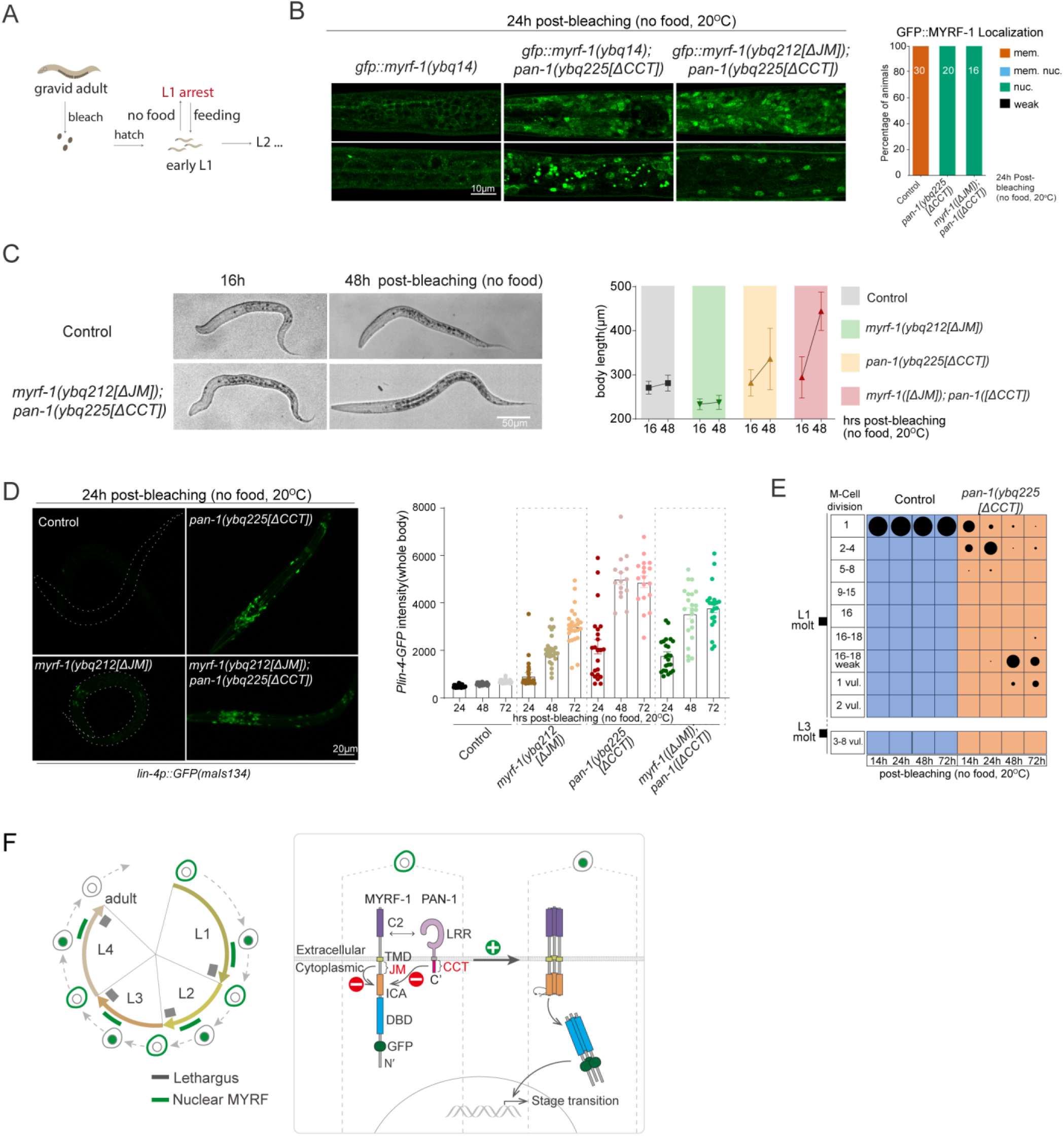
Disruption of L1 arrest by MYRF-1[ΔJM] and PAN-1[ΔCCT]. (A) Schematic of the L1 diapause assay. Embryos hatched on food-less plates enter diapause and resume development upon refeeding. (B) GFP::MYRF-1 localization after 24 h starvation. Controls show membrane localization; *pan-1[ΔCCT]* and *myrf-1[ΔJM]; pan-1[ΔCCT]* mutants show predominant nuclear localization. Right: quantification of GFP::MYRF-1 localization. (C) Larvae at 16 and 48 h starvation. *myrf-1[ΔJM]; pan-1[ΔCCT]* mutants are longer than controls. Right: body length quantification (mean ± SD). (D) *lin-4p::GFP* expression at 24 h starvation; it increases in mutants. Right: quantification of reporter intensity at 24, 48, and 72 h starvation (mean ± SEM). (E) M-cell lineage progression in control and *pan-1[ΔCCT]* mutants at 14, 24, 48, and 72 h starvation; *hlh-8p::gfp* used for staging. (F) Summary model. Left: MYRF cleavage peaks oscillate during larval stages. Right: cleavage is suppressed by the MYRF JM region and PAN-1 CCT; inhibition is likely relieved by a systemic cue enabling trimerization and self-cleavage.

When this cleavage inhibition is removed, L1 arrest is disrupted. Body length gradually increased in *pan-1[ΔCCT]* mutants and even more prominently in *gfp::myrf-1[ΔJM]; pan-1[ΔCCT]* double mutants. By 48 hours, the double mutants reached lengths comparable to L2 larvae, while wild-type L1s remained unchanged (Figure 6C). This growth indicates slow but significant developmental progression in the absence of food. Moreover, *lin-4* transcription—normally inactive under starvation—was prematurely activated in both *gfp::myrf-1[ΔJM]* and *pan-1[ΔCCT]* single mutants, with stronger induction in *pan-1[ΔCCT]* (Figure 6D). M-cell divisions also occurred in *pan-1[ΔCCT]* mutants, reaching late-L1–like patterns (Figure 6E; Figure 6—figure supplement 1A), whereas *gfp::myrf-1[ΔJM]* mutants showed only minimal precocious division (Figure 6—figure supplement 1B). Together, these findings demonstrate that loss of MYRF-1 cleavage inhibition permits developmental progression during starvation, at least through the L1 stage.

Larvae enter L1 diapause to prolong survival under starvation and resume normal development when food becomes available. Relief of MYRF cleavage inhibition impaired the animals’ ability to recover. After 48 hours without food, both *pan-1[ΔCCT]* single and *gfp::myrf-1[ΔJM]; pan-1[ΔCCT]* double mutants showed poor developmental recovery upon refeeding, with higher rates of arrest compared to continuously fed animals (Figure 6—figure supplement 1C). This suggests that proper regulation of MYRF-1 cleavage is crucial not only for arrest but also for enabling robust recovery when conditions improve.

## Discussion

We previously demonstrated that cleavage is essential for MYRF-1 function in *C. elegans*, but the dynamics and regulatory significance of this cleavage remain undetermined. Here, we show that MYRF-1 cleavage is under tight temporal and oscillates across larval stages—peaking in mid-to-late phases and being suppressed during molts (Figure 6F). This oscillatory pattern aligns with MYRF-1’s role in promoting stage transitions, particularly late-L1 events such as *lin-4* activation and DD neuron remodeling.

We identified two inhibitory elements—an intrinsic JM region within MYRF-1 and an extrinsic CCT domain from its binding partner PAN-1—that suppress MYRF-1 cleavage, with the PAN-1 CCT serving as the dominant control (Figure 6F). Genetic removal of these inhibitory elements disrupts cleavage timing, triggers precocious activation of downstream events, and ultimately causes developmental arrest or lethality. These findings establish MYRF-1 cleavage as a key regulatory node in the *C. elegans* developmental timing network.

Our data suggest that MYRF-1 drives progression through larval stages, particularly at late L1. Consistent with this, *myrf-1* null mutants arrest during molting and fail to activate *lin-4* or execute DD wiring^[7,9,15]^. In contrast, earlier L1 events such as P-cell and M-cell divisions are less affected^[7]^. However, the roles of MYRF-1 in later stages (L2–L4) remain unclear due to early arrest and may require temporally controlled perturbation. Importantly, *myrf-1* is among over 3,700 genes that exhibit transcript-level oscillation^[16,22]^, and it contributes to the regulation of this network^[23]^. How MYRF-1 interacts with other key oscillatory factors—such as GRH-1, BLMP-1, NHR-23, NHR-25, and BED-3^[23,24]^—warrants further investigation.

MYRF-1 may integrate nutritional cues with developmental timing. Under starvation, MYRF-1 cleavage is blocked, aligning with diapause entry. Disrupting cleavage inhibition (e.g., in *pan-1[ΔCCT]* or *myrf-1[ΔJM]* mutants) leads to premature *lin-4* activation and partial bypass of nutrient-induced arrest at early L3 and L4. Notably, M-cell division occurs in *pan-1[ΔCCT]* mutants during L1 diapause despite being dispensable for *myrf-1* function under normal conditions^[7]^. This implies that nuclear MYRF-1 might trigger broader stage transitions that feed back to earlier events, reflecting a hierarchical control logic in temporal regulation. However, relieving cleavage inhibition also compromises recovery from arrest, underscoring the importance of precise regulation in fluctuating environments.

Mechanistically, both the JM and PAN-1 CCT domains are positioned to interact with the ICA domain, which mediates MYRF trimerization and autocleavage. The JM region appears to confer cis-inhibition, likely by sterically or conformationally hindering trimer formation during or shortly after MYRF-1 translation, while PAN-1 CCT may reinforce cleavage suppression once MYRF-1 is membrane-inserted in ER. The factor(s) responsible for relieving this inhibition remain unknown but may involve extracellular ligands or post-translational modifications that alter MYRF-PAN-1 interactions or expose the ICA domain. Identifying such factors will be key to understanding how temporal signals converge on MYRF-1.

Finally, the developmental regulation and function of mammalian MYRF remain incompletely defined beyond its well-known role in oligodendrocyte myelination^[1]^. Our findings in *C. elegans* show that membrane-tethered MYRF can be tightly regulated through interacting partners such as PAN-1. While no PAN-1 ortholog is known in mammals, TMEM98, which is conserved in *C. elegans* and mammals, was recently shown to bind to MYRF and inhibit its cleavage in mouse, suggesting that a conserved regulatory logic may exist^[25,26]^. The principles uncovered here may thus illuminate broader mechanisms by which cleavage-activated transcription factors coordinate development with internal and external cues.

## Materials and Methods

### Animals

Wild-type *C. elegans* used in this study were of the Bristol N2 strain. Worms were cultured on NGM plates following standard procedures. Unless otherwise specified, animals were maintained at 20°C for assays requiring specific developmental stages. All animals analyzed in this study were hermaphrodites.

### Allele and Strain Nomenclature

All alleles generated in the Y.B.Q. lab are designated with the "ybq" prefix, and all strains are referred to as "BLW" strains. The designation "Ex" indicates extrachromosomal array transgenes, while "Is" denotes integrated transgenes, and "Si" represents single-copy integrated transgenes. "syb" alleles (in "PHX" strains) are produced via genomic editing using the CRISPR-Cas9 technique. These "syb" alleles were designed by Y.B.Q. and generated by SunyBiotech (Fuzhou, China).

### MYRF-1 ΔJM allele by CRISPR-Cas9 editing

*gfp::myrf-1(ybq212[ΔJM(K601-C650)])* was generated by CRISPR-Cas9 editing^[27,28]^. It was made in the background of strain BLW2171 (*GFP::MYRF-1::3xFlag(ybq14[gfp::A172]) II; oxTi1128 [mex-5p::Cas9(+smu-2 introns)::tbb-2 3’UTR + hsp-16.41p::Cre::tbb-2 3’UTR + myo-2p::2xNLS::cyOFP::let-858 3’UTR + lox2272]) I*). The vectors expressing gRNAs and repair templates vectors were together microinjected. Successfully edited alleles were identified through PCR screening of individual F1 progeny resulting from the microinjections. The Cas9 transgene (*oxTi1128*) was removed through subsequent outcrossing or during crosses with balancer strains. The resulted *myrf-1 ΔJM* allele removes the JM(K601-C650) region of MYRF-1 coding sequence. For simplicity, in this paper, this allele that also carry the endogenous GFP tag (*ybq14*) are referred to as *gfp::myrf-1(ybq212[ΔJM])*, *myrf-1(ybq212[ΔJM])*, *myrf-1(ybq212)*, or *myrf-1[ΔJM]*.

gRNA targets are:

Sg1: AAAGAAGCTCGCTAGCTCAATGG;
Sg2: GCTTCTCTTTGGTTGTGACGTGG;
Sg3: ATAAAACAGGCACAATCCTGTGG;
Sg4: TGTGGAAGCCGACTTAGTCAAGG

*gfp::myrf-1(ybq14)*: …AGACGATTGAATGAATACGCAGTCCGAAAG-(150bp)-GGAAGCCGACTTAGTCAAGGAACAGTTGTC

*gfp::myrf-1(ybq14 ybq212[ΔJM])*: …AGACGATTGAATGAATACGCAGTCCGAAAG-(delete 150bp)-GGAAGCCGACTTAGTCAAGGAACAGTTGTC…

### MYRF-1 S623A allele by CRISPR-Cas9 editing

The alleles myrf-1(ybq215[S623A]) were generated in the background of strain BLW2171 (myrf-1(ybq14[gfp::myrf-1::3xflag]) (GFP is at A172) II; oxTi1128 [mex-5p::Cas9(+smu-2 introns)::tbb-2 3’UTR + hsp-16.41p::Cre::tbb-2 3’UTR + myo-2p::2xNLS::cyOFP::let-858 3’UTR + lox2272]) I), following the procedure described in “Generate MYRF-1 ΔJM allele by CRISPR-Cas9 editing”. For simplicity, in this paper, this gene-edited allele that also carries the endogenous GFP tag (ybq14) are referred to as gfp::myrf-1(ybq215[S623A]).

The myrf-1(ybq215[S623A]) allele was generated in the BLW2171 strain background (genotype: myrf-1(ybq14[gfp::myrf-1::3xflag]) (GFP inserted at A172) II; oxTi1128 [mex-5p::Cas9(+smu-2 introns)::tbb-2 3’UTR + hsp-16.41p::Cre::tbb-2 3’UTR + myo-2p::2xNLS::cyOFP::let-858 3’UTR + lox2272]) I) using the CRISPR-Cas9 editing protocol described in “MYRF-1 ΔJM allele by CRISPR-Cas9 editing”. For brevity, this gene-edited allele—which retains the endogenous GFP tag from the parental ybq14 allele— is hereafter referred to as gfp::myrf-1(ybq215[S623A]).

gRNA targets are:

Sg1: AAAGAAGCTCGCTAGCTCAATGG;
Sg2: GCTTCTCTTTGGTTGTGACGTGG;

gfp::myrf-1(ybq14):

…AGACGATTGAATGAATACGCAGTCCGAAAG - (insertion point) - AAGCTCGCTAGCTCAATGGCATCTAACCTCAACGGAGACAACAAGAGTCTCTCATACTCAAG - (insertion point) - ATGCTCACTCACATCAACTGCAACCAATGCCACGTCACAACCAAAGAGAAGCAGGAAGCAT..
gfp::myrf-1(ybq215[S623A]): …AGACGATTGAATGAATACGCAGTCCGAAAG - (gtataaggttaaacaaaagtttatttttcaaaaacgttgttttcag) - AAGCTCGCTAGCTCAATGGCATCTAACCTCAACGGAGACAACAAGAGTCTCTCATACTCAAG
- (gtaaatttctttttttttcagcaaaaattctaatgtttcattttttag) - ATGCgCACTCACATCAACTGCAACCAATGCtACGTCACAACCAAAGAGAAGCAGGAAGCAT…

### PAN-1 ΔCCT allele by CRISPR-Cas9 editing

*pan-1(ybq225[ΔCCT(R542-F591)])* was generated in the background of strain BLW2171 (*GFP::MYRF-1::3xFlag(ybq14) II; oxTi1128 [mex-5p::Cas9(+smu-2 introns)::tbb-2 3’UTR + hsp-16.41p::Cre::tbb-2 3’UTR + myo-2p::2xNLS::cyOFP::let-858 3’UTR + lox2272]) I*), following the procedure described in “MYRF-1 ΔJM allele by CRISPR-Cas9 editing”. The resulted allele removes the CCT(R542R-F591) region of PAN-1 coding sequence.

gRNA targets are:

Sg1: TGGAATCGTACAGTCTTCTTCGG;
Sg2: aggaaaaactaatatatacCTGG
*pan-1*(wt): …ATTATTGCTATGCTTTATTTCAAGGATGCT-(207bp)-CAGAATTTTTAAagaagaaaaatttcataa…
*pan-1(ybq225[ΔCCT])*: …ATTATTGCTATGCTTTATTTCAAGGATGCT-(delete 207bp)-CAGAATTTTTAAagaagaaaaatttcataa…

### C-terminally inserted GFP allele of *myrf-1* by CRISPR-Cas9 editing

*myrf-1(syb3705[GFP::G651])* was generated in the background of wild-type (N2). GFP is inserted before G651 of the MYRF-1 coding sequence.

WT: …CAGGAAGCATCGTGCCATAAAACAGGCACAATCCTGT-(insertion point)-GGAAGCCGACTTAGTCAAGGAACAGTTGTCACCTTGG…

*myrf-1(syb3705)*: …CAGGAAGCATCGTGCCATAAAACAGGCACAATCCTGT-GGTGGCGGAGGTTCT(GGGGS linker)-mEGFP(A206K)-GGAAGCCGACTTAGCCAAGGAACAGTTGTCACCTTGG…

### Generation of Extrachromosomal Array and Single-Copy Insertion Transgene Alleles

Overexpression of NLS::GFP::MYRF-1[G477-C’]: The vector pQA1815 [Prpl-28::NLS::GFP::myrf-1[G477-C’] with miniMos] was injected into RW10434 strain at a concentration of 10 ng/µl to generate overexpressing lines.

Overexpression of NLS::GFP::MYRF-1[G477-C’, ΔJM(601-650)]: The vector pQA1877 [Prpl-28::NLS::GFP::myrf-1[G477-C’, ΔJM(601-650)] with miniMos] was injected into RW10434 strain at 10 ng/µl for overexpression studies.

Single-Copy Transgene of *Pscm-tagRFP::PH (ybqSi296)*: The vector pQA2337 [Pscm-tagRFP::PH domain] with miniMos] was injected into BLW1827[GFP::MYRF-1::3xFlag(ybq14) II; glo-1(zu391)] at 50 ng/µl along with standard miniMos system components.

### Single-copy integrated transgene allele

The procedure for generating single-copy integrated transgenes using miniMos technology^[29]^. The injection mixture contained 10 ng/µl pGH8 [rab-3p::mCherry], 2.5 ng/µl pCFJ90 [myo-2p::mCherry], 50 ng/µl pCFJ601 [eft-3p::Mos1-transposase], and 50 ng/µl miniMos plasmid harboring the target gene. The mixture was injected into the gonads of N2 animals. After injection, the nematodes were transferred onto NGM medium and incubated at 25°C for approximately 48 hours. Subsequently, 500 µl of 25 mg/mL G418 solution was added to each plate to screen for nematodes carrying the target transgene. After 7-10 days, healthy nematodes without mCherry co-markers were selected from the medium where all the food was consumed. Candidate single-copy integrators were grown on G418-containing plates and analyzed for target protein expression. Homozygous nematodes carrying a single-copy transgene were identified by their 100% resistance to G418 toxicity and expression of the target protein.

### Microscopic Analyses and Quantification

Live animals were washed in M9, anesthetized in a small drop of 0.5-1% 1-Phenoxy-2-propanol in ddH2O on coverslip, and mounted on 3% agarose gel pads. The animals were examined using 20x, 40x, or 60x oil objectives on an OLYMPUS BX63 microscope. Wide-field DIC or fluorescence images (either single-plane or Z-focal stacks) were acquired using a sCMOS camera (PrimΣ Photometrics model 2) mounted on the OLYMPUS BX63, controlled by CellSens Dimension software (Olympus). Images of live animals were also acquired using a Zeiss LSM980 with Airyscan 2, with the optical slice thickness typically set to 0.8 µm.

To quantify the patterns of GFP::MYRF-1, we acquired images of stage-synchronized animals using either wide-field sCMOS camera or Zeiss Airyscan, as described above. Consistent parameters were used, including identical excitation light power, objectives, exposure duration, and display settings. The images were analyzed, and GFP::MYRF-1 patterns were categorized based on the consistency of signal localization in specific subcellular regions throughout the animal. "Weak or unclear signal" refers to either no detectable signals or weak, inconsistent signals throughout the animal body.

For fluorescence intensity measurements of GFP::MYRF-1, the head region was selected. GFP::MYRF-1 signals are relatively weak, and choice of head region is to avoid gut autofluorescence, which can significantly interfere with measurements in the mid-body. Z-stacks were maximally projected into single images using ImageJ. The entire head area was selected as the region of interest (ROI), and the mean intensity of the ROI was measured and presented as Mean ± SEM. Statistical significance was determined using a t-test (*p < 0.05; **p < 0.01; ***p < 0.001; ****p < 0.0001).

For quantifying fluorescence intensity of *lin-4p::GFP(maIs134)*, Z-stack images were maximally projected into single images using ImageJ, with the entire body area selected as the ROI. The mean intensity of the ROI was measured and presented as Mean ± SEM. Statistical significance was determined using a t-test (*p < 0.05; **p < 0.01; ***p < 0.001; ****p < 0.0001).

To quantify the whole body *lin-4p::nls::mScarlet(umn84)* transcriptional reporter fluorescence intensity, the z-stack was maximally projected to produce a single image in ImageJ. The whole animal area was selected as ROI. The intensity of the ROI was measured and presented as Mean ± SEM (t-test, *, p<0.05; **, p<0.01; ***, p<0.001; ****, p<0.0001).

For quantifying M cell divisions labeled by *ayIs6[hlh-8p::GFP]*, Z-stack images were acquired using a 20x objective lens on an OLYMPUS BX63 microscope. The number of ayIs6-labeled cells was visually identified and recorded. Graphs were generated using Microsoft Excel.

For quantifying seam cell divisions labeled by *heIs63 [wrt-2p::GFP::PH + wrt-2p::GFP::H2B + lin-48p::mCherry]* or *ybqSi296(Pscm::tagRFP::PH)*, Z-stack images were acquired using a 20x objective lens on an OLYMPUS BX63 microscope. The cell morphology and number of marker-labeled V1-V6 seam cells was visually identified and recorded.

### Collecting worms in lethargus

Synchronized early L1 worms were cultured on OP50-seeded NGM plates and maintained at 20°C. At specific developmental timepoints—16–17 hours post-hatch for L1 lethargus, 27–28 hours for L2 lethargus, 36–37 hours for L3 lethargus, and 46–47 hours for L4 lethargus—worms undergoing lethargus were identified and transferred onto fresh OP50-seeded NGM plates for subsequent imaging. Worms in lethargus were identified as those with reduced movement and cessation of pharyngeal pumping for at least 20 seconds.

### L1 worm synchronization by bleaching and L1-diapause Induction

Gravid adults were washed off the plate and rinsed three times with water by sedimentation at room temperature. The loose worm pellet was then mixed with freshly made bleach buffer (1 M NaOH : CLOROX Multipurpose bleach in a 5:2 ratio). Disintegration of the worm bodies was monitored under a stereoscope. Once disintegration began, the tube was quickly spun down, and the remaining worms were rinsed three times in M9 buffer before being transferred onto non-food plates. The worms were kept on the non-food plates for a designated number of hours before analysis.

### Food Deprivation Assay

Plate-based deprivation assay for late L1 plus 8h starvation: Gravid adults were lysed using bleach solution to collect eggs, which were washed three times with M9 buffer and transferred to food-free NGM plates allowing hatching for no longer than 20 hours. Synchronized early L1 larvae were cultured on OP50-seeded NGM plates for the indicated durations post-hatching. Approximately 200 larvae were then washed in M9 buffer to remove residual bacteria (centrifugation at 250 × g for 1 minute, repeated for 5 times), and transferred to either food-free or OP50-containing NGM plates for an additional 8 hours of culture.

M9 buffer-based deprivation assay (late L1, L2, L3, L4 plus 24h/48h starvation): Gravid adults were bleached and embryos transferred to 8 or 15 mL glass vials (loosely capped) containing 1 mL M9 buffer. Vials were shaken at 90 rpm at 20°C for no more than 20 hours to allow hatching. Synchronized early L1 larvae were then cultured on OP50 plates until the desired developmental stages. Worms were washed five times in M9 buffer to remove bacteria and transferred to glass vials containing 1 mL fresh M9. Vials were shaken at 90 rpm at 20°C for the indicated durations.

### Quantification of Animal Length and Body Area

Live animals were anesthetized with 0.5% 1-Phenoxy-2-propanol in ddH2O and mounted on a 3% agarose gel pad. Differential Interference Contrast (DIC) images were acquired using an OLYMPUS BX63 microscope with a 10x objective. The length of each animal was measured using the polyline tool in the CellSens Dimension software (Olympus). Body area (in pixels) was measured using ImageJ software. The collected data were then analyzed using GraphPad Prism.

### Brood Size Analysis

Synchronized early L1 worms were cultured on OP50 plates for 48 hours. Individual worms were then transferred to fresh OP50 plates, and transfers were repeated every 12 hours for 5 days (n > 30 per group). The progeny on each plate were counted and summed for each individual.

### Survival Analysis

Approximately 200 L4-stage worms were transferred and distributed across four OP50-seeded NGM plates, then cultured at 20°C. Animals were transferred to fresh plates daily until the end of the reproductive period. Counting continued until all *myrf-1[ΔJM]* mutants had died.

### Western Blot Analysis

BLW889 [GFP::MYRF-1::3xFlag(ybq14)] animals were cultured to the desired developmental stage, collected, and washed twice with M9 buffer. After centrifugation and sedimentation on ice, the volume of the worm pellet was recorded. Worm pellets were resuspended in an equal volume of pre-chilled 2× lysis buffer (100 mM Tris-HCl pH 7.5, 300 mM NaCl, 2% NP-40, 1% sodium deoxycholate, 2% SDS, 2 mM EDTA, and 2× Roche complete EDTA-free protease inhibitor cocktail), and lysed by vigorous shaking on ice for 45 minutes. Lysates were denatured at 70°C for 15 minutes, then centrifuged at 12,000 rpm for 10 minutes at 4°C. The resulting supernatants were subjected to SDS-PAGE followed by western blotting. The following antibodies were used: Mouse anti-GFP (Abmart, M200004F; 1:2000); Mouse anti-β-actin, HRP-conjugated (Beyotime, AF2815; 1:2000); Goat anti-mouse IgG (H&L), HRP-conjugated (Genscript, A00160; 1:10,000).

## Key Resources Table

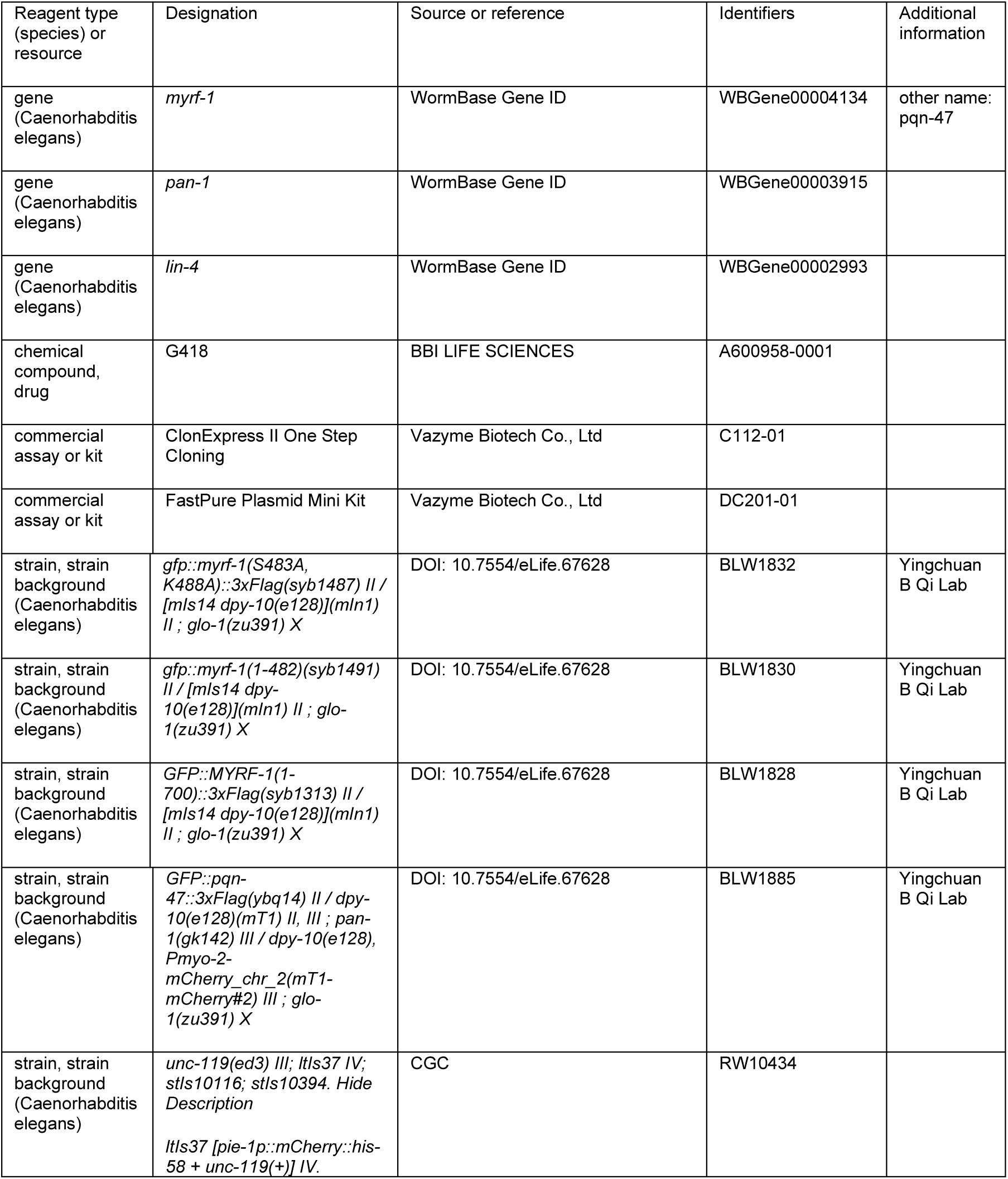
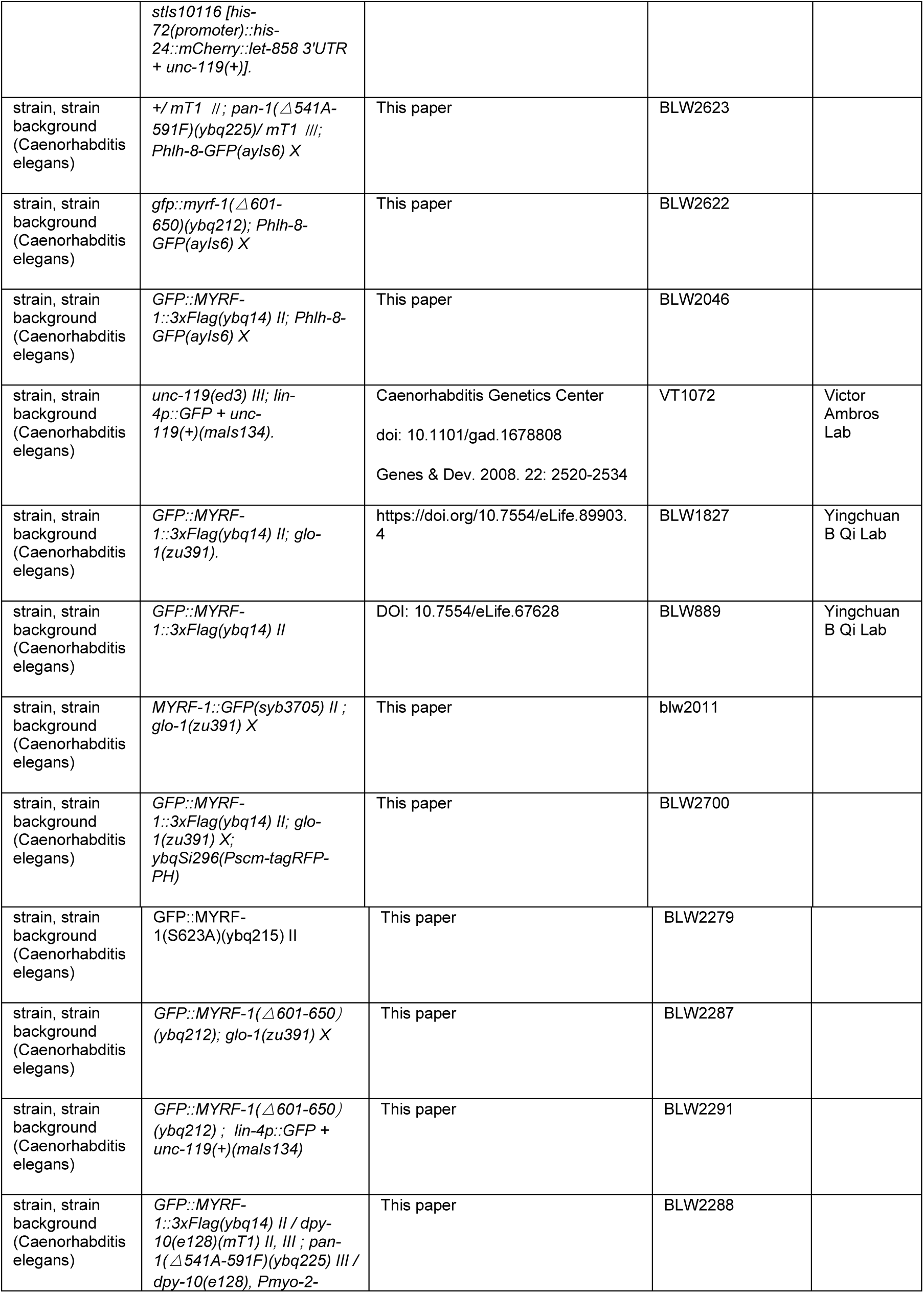
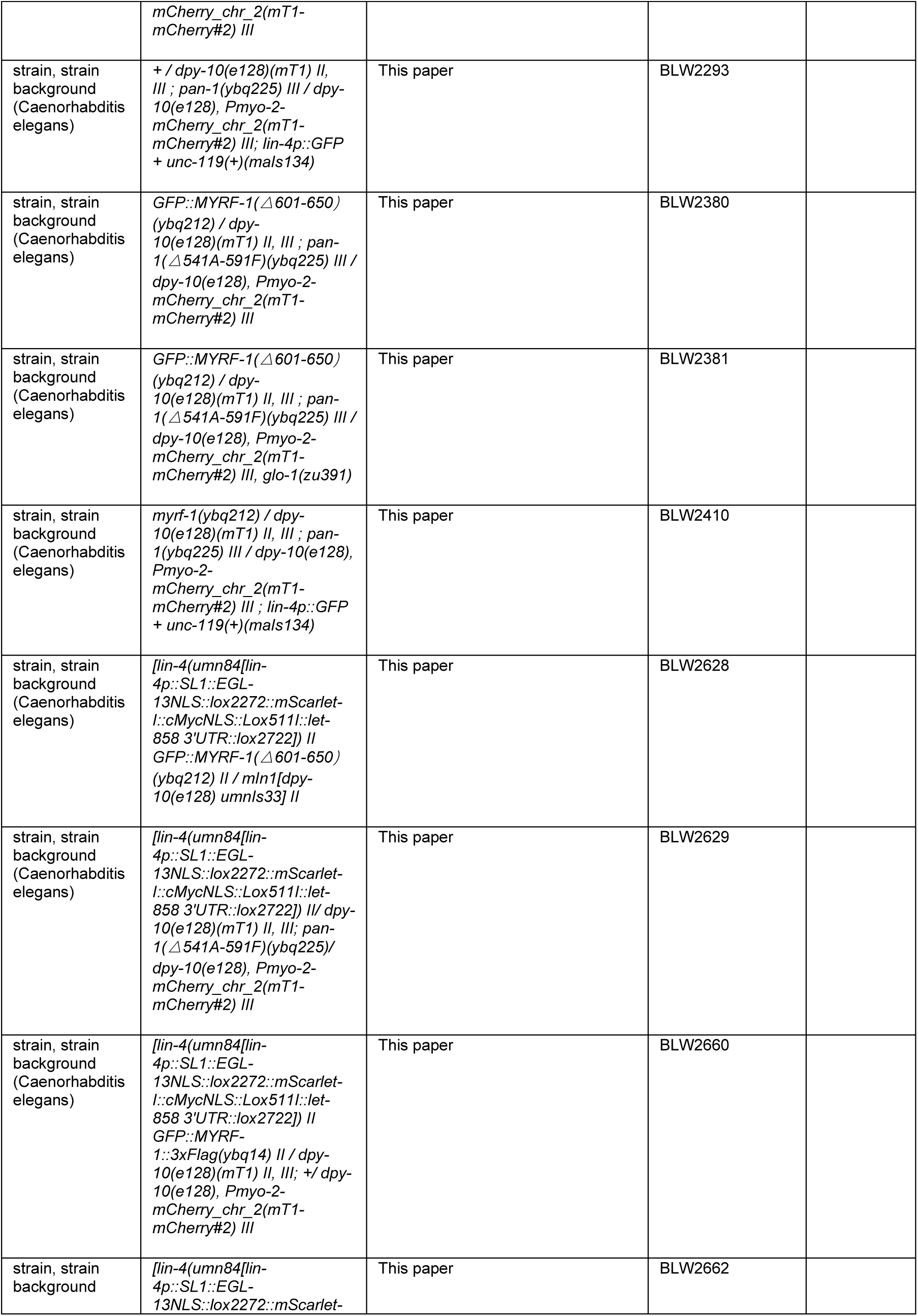
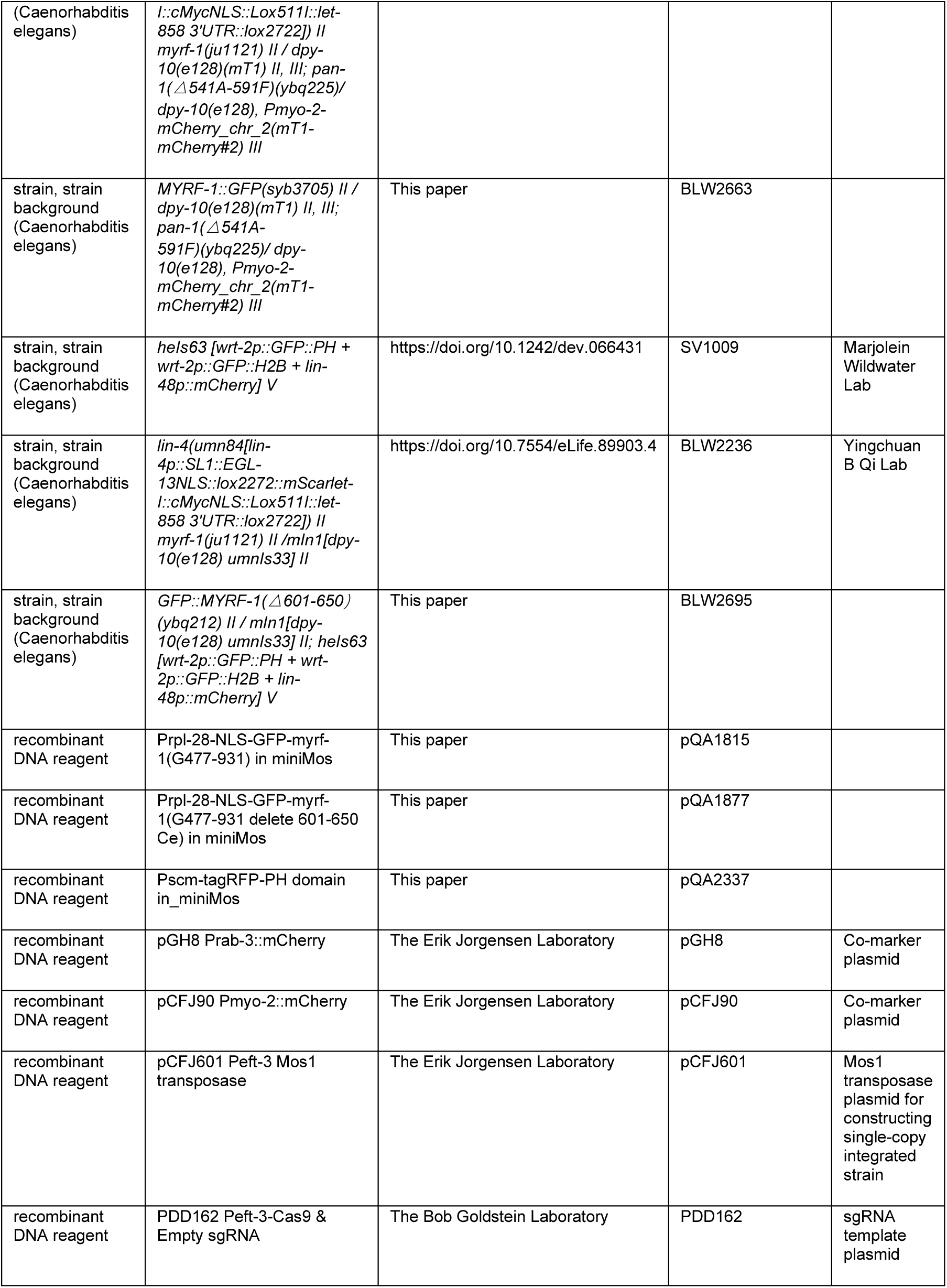

## Acknowledgments

We thank Qian Bian, Di Chen, Yan Zou, Huanhu Zhu, and Gaofeng Fan for their valuable discussions. We also thank the Molecular and Cell Biology Core Facility (MCBCF) and the Molecular Imaging Core Facility (MICF) at the School of Life Science and Technology, ShanghaiTech University for providing technical support. Some strains were provided by the CGC, which is funded by NIH Office of Research Infrastructure Programs (P40 OD010440). This work was supported by the National Key R&D Program of China (Grant Nos. 2022YFC3400501 and 2022YFC3400500) to F.B., the start-up fund from ShanghaiTech University to Y.B.Q., and the National Science Foundation of China (Grant No. 32370601) to Y.B.Q.

---

**Figure 1—figure supplement 1.**
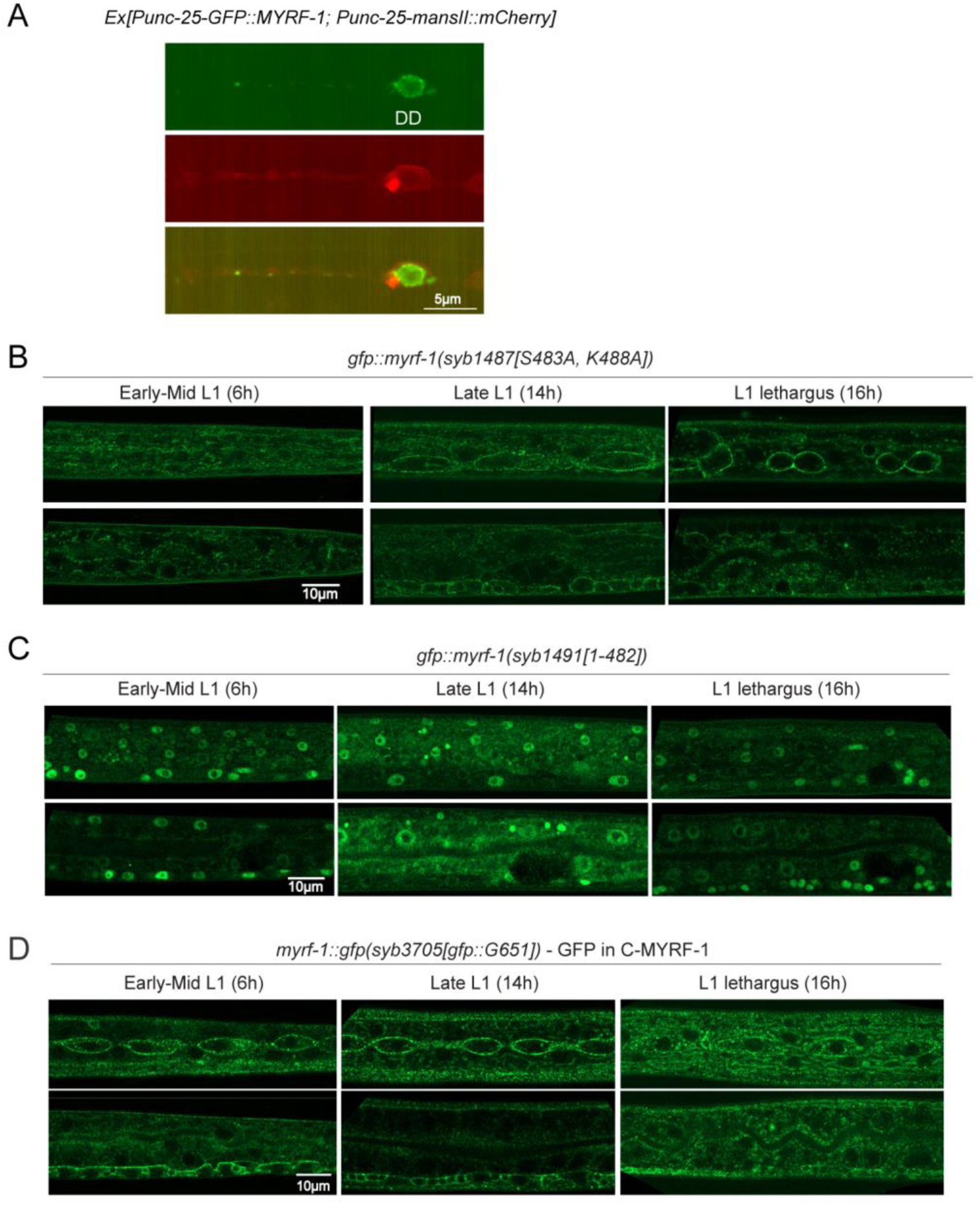
Cleavage pattern and localization dynamics of MYRF-1. (A) Overexpressed GFP::MYRF-1 in DD neurons localizes to the cytoplasm and overlaps with the Golgi marker manII::mCherry. (B) GFP::MYRF-1(*syb1487[S483A, K488A]*) shows membrane localization in the trunk (top: epidermis, seam; bottom: intestine, VNC) across L1. Complements head-region images in Figure 1E. (C) GFP::MYRF-1(*syb1491[1–482]*) shows nuclear localization in trunk across L1. Complements Figure 1F. (D) C-terminal GFP fusion in *myrf-1::gfp(syb3705)* localizes to the cell membrane in the trunk across L1. Complements Figure 1G. Right: Schematic shows GFP insertion at Gly651.

**Figure 2—figure supplement 1.**
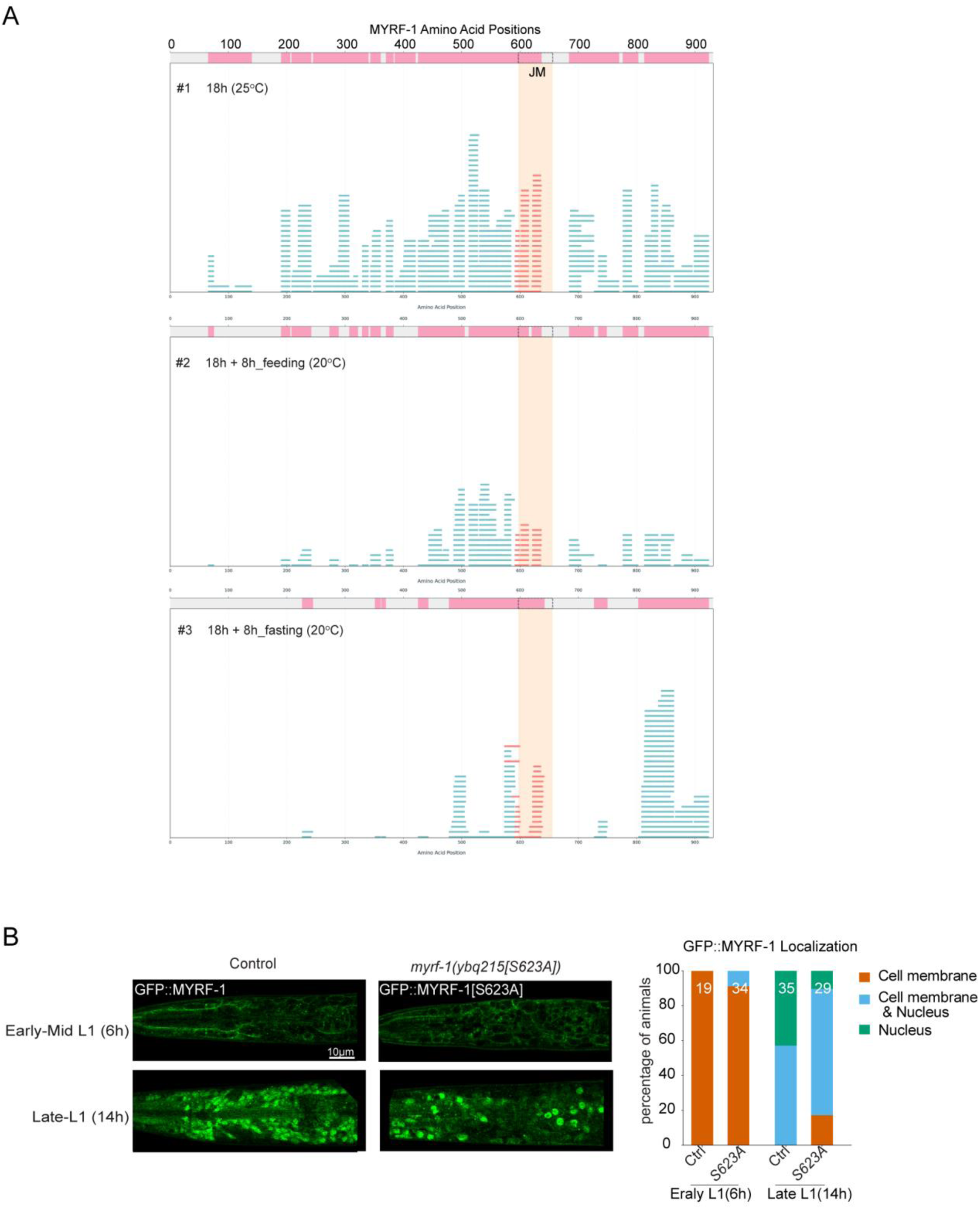
IP–MS identifies a potential regulatory region in MYRF-1. (A) Peptide coverage of MYRF-1 across three IP–MS samples. The juxtamembrane (JM) region is highlighted in peach. Each blue line represents an identified peptide. (B) Representative localization of GFP::MYRF-1(ybq215[S623A]) at 6 h and 14 h post-hatching. Right: Quantification of subcellular localization.

**Figure 3—figure supplement 1.**
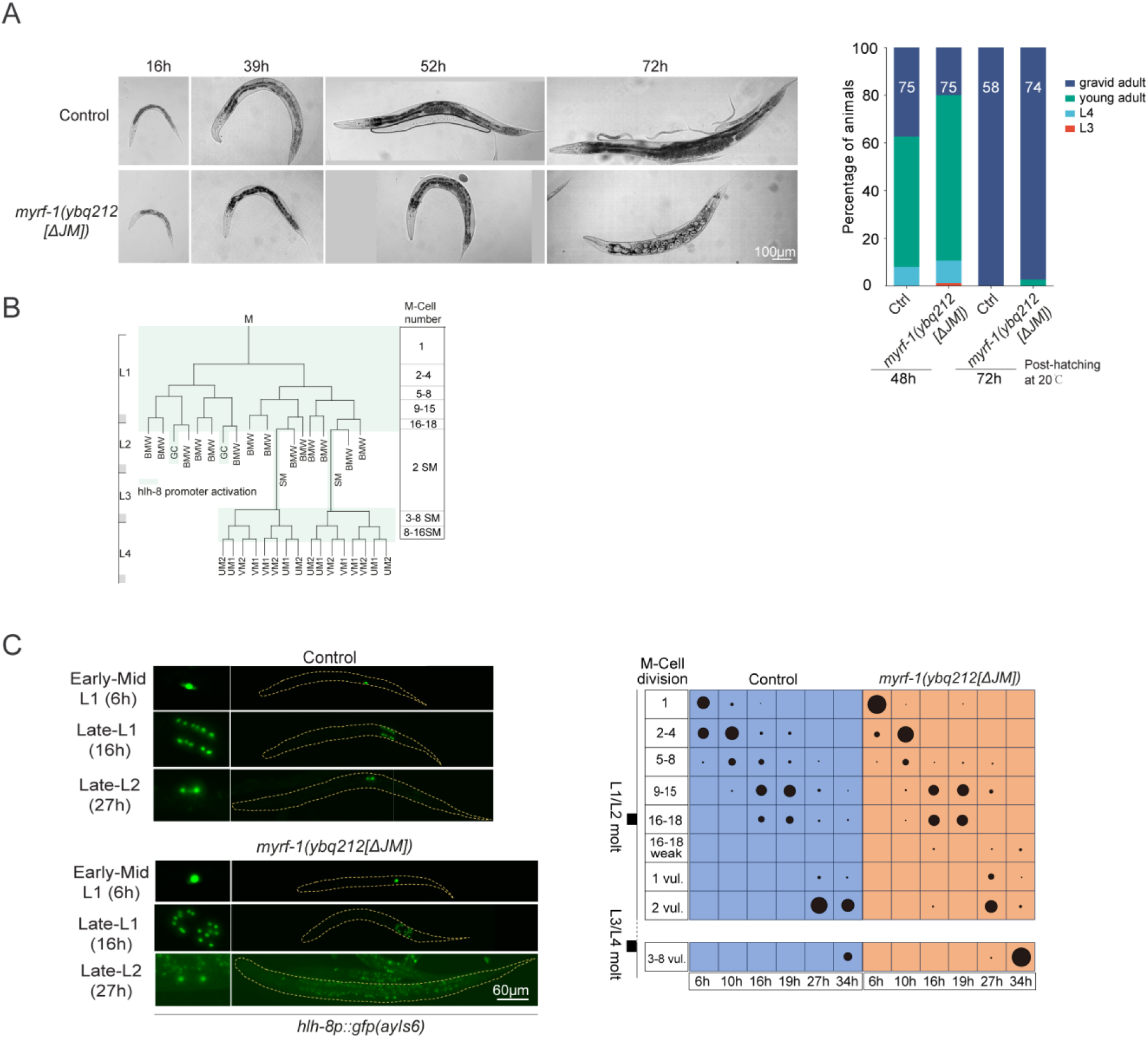
Characterization of *myrf-1(ybq212[ΔJM])* mutants. (A) Representative *myrf-1[ΔJM]* mutants during larval development. Mutants are shorter than controls; adults exhibit bagging. Right: Developmental progression of *gfp::myrf-1(ybq14)* controls and *myrf-1[ΔJM]* mutants at 48 h and 72 h post-hatching. (B) Schematic of M-cell lineage: hlh-8p::GFP marks undifferentiated M-cell progeny; expression is lost upon muscle differentiation. (C) hlh-8p::GFP expression in M-cell lineages of control and *myrf-1[ΔJM]* mutants at 6, 16, and 27 h. Right: quantification of M-cell division stages.

**Figure 5—figure supplement 1.**
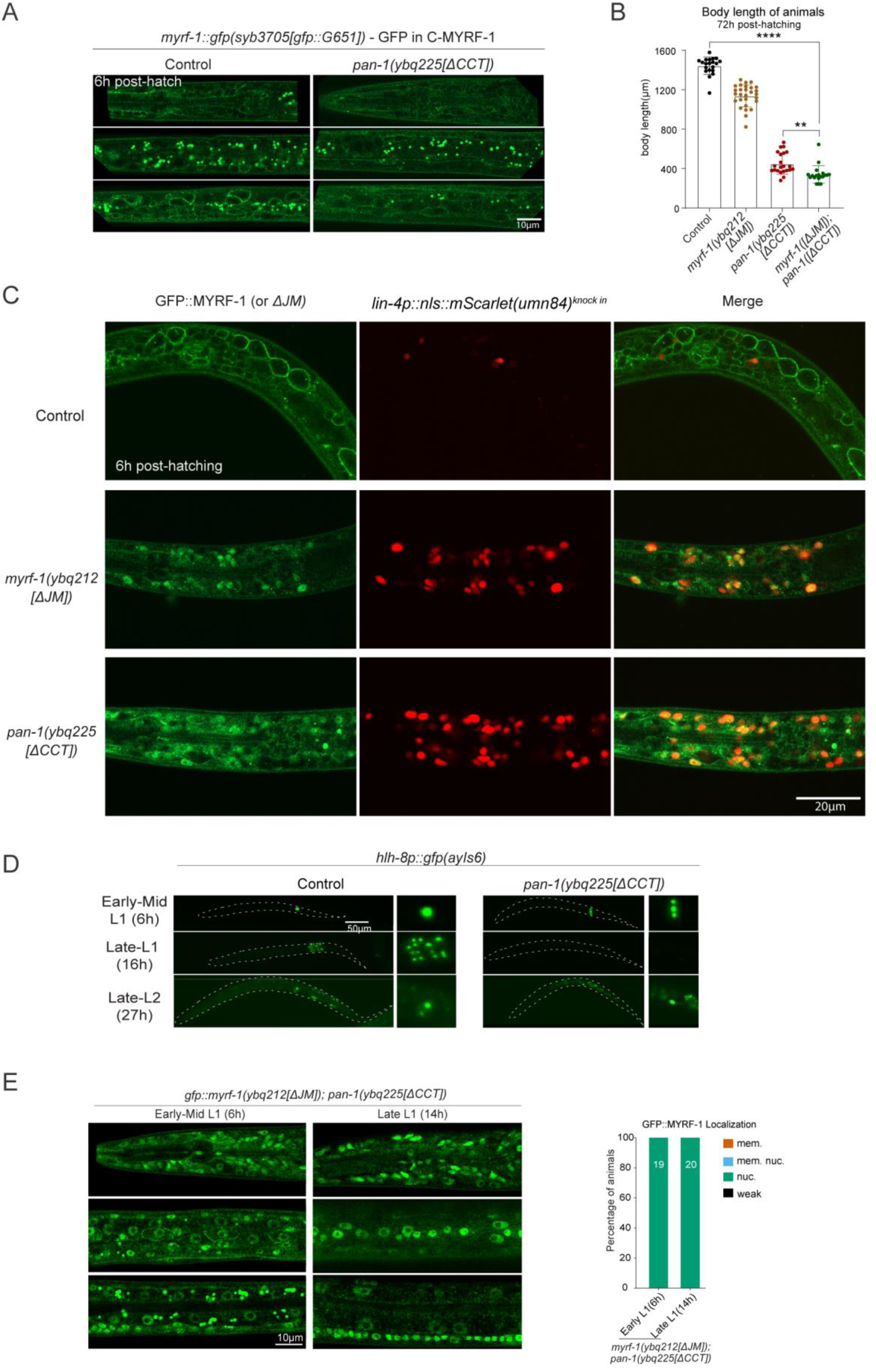
Characterization of *pan-1(ybq225[ΔCCT])* mutants. (A) In *pan-1[ΔCCT]* mutants, MYRF-1(*syb3705[GFP::G651]*) remains membrane-localized at 6 h. Sagittal sections show mid-head, lateral trunk, and mid-trunk regions. Complements Figure 5D. (B) Body length measurements of control, *myrf-1[ΔJM]*, *pan-1[ΔCCT]*, and *myrf-1[ΔJM]; pan-1[ΔCCT]* animals post-hatching (mean ± SD; **p < 0.01; ****p < 0.0001; t-test). (C) Co-labeling of GFP::MYRF-1 (*ybq14*), *gfp::myrf-1[ΔJM]*, or GFP::MYRF-1 in *pan-1[ΔCCT]* with endogenous *lin-4* reporter (*umn84*) at 6 h. *lin-4* is not activated in wild type, but precociously activated in both mutants, colocalizing with nuclear MYRF-1. (D) hlh-8p::GFP(*ayIs6*) marking M-cell divisions in control and *pan-1[ΔCCT]* mutants at 6, 16, and 27 h. (E) In *myrf-1[ΔJM]; pan-1[ΔCCT]* double mutants, GFP is predominantly nuclear. Right: quantification of GFP localization.

**Figure 6—figure supplement 1.**
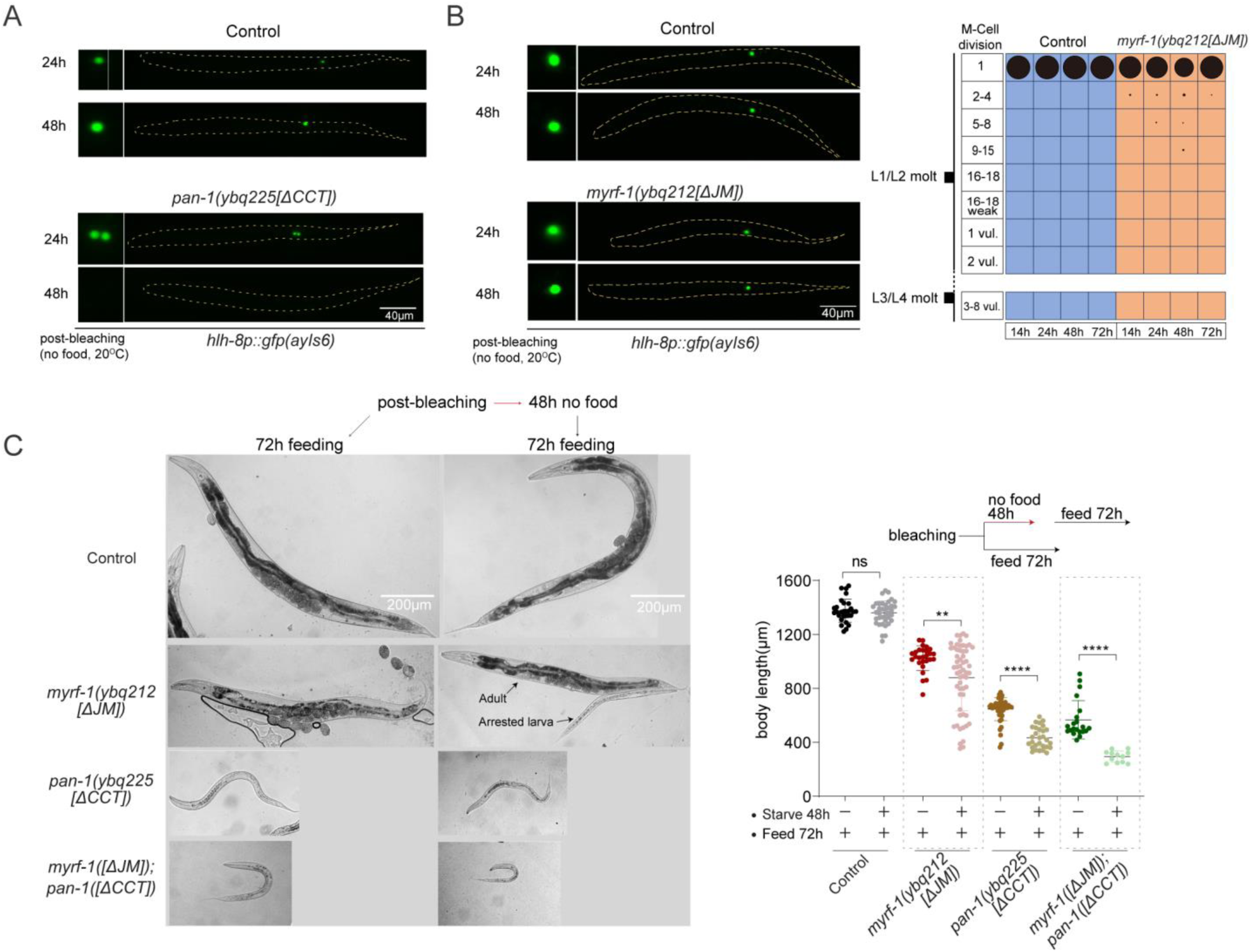
Disruption of L1 diapause by relieving MYRF-1 cleavage inhibition. (A) Representative *pan-1[ΔCCT]* animals under early L1 starvation (24 h and 48 h), with M cells labeled by hlh-8p::GFP(ayIs6). M-cell divisions occur during diapause in mutants but not in controls. By 48 h, mutants show 16–18 M-lineage cells with weak hlh-8p::GFP expression. Complements Figure 6E. (B) *myrf-1[ΔJM]* mutants under early L1 starvation for 24 h and 48 h. M-cell divisions are not advanced. Right: Quantification of M-lineage patterns. (C) Recovery ability after 48 h of early L1 starvation. Animals were either starved post-bleaching followed by 72h recovery or cultured with food continuously. Right: Body length measurements (mean ± SD; >20 animals/group).

